# Prednisolone rescues Duchenne Muscular Dystrophy phenotypes in human pluripotent stem cells-derived skeletal muscle *in vitro*

**DOI:** 10.1101/2020.10.29.360826

**Authors:** Ziad Al Tanoury, John F. Zimmermann, Jyoti Rao, Daniel Sieiro, Harry McNamara, Thomas Cherrier, Aurore Hick, Fanny Bousson, Charlotte Fugier, Fabio Marchiano, Bianca Habermann, Jérome Chal, Alexander P. Nesmith, Svetlana Gapon, Erica Wagner, Rhonda Bassel-Duby, Eric Olson, Adam E. Cohen, Kevin Kit Parker, Olivier Pourquié

**Affiliations:** Institut de Génétique et de Biologie Moléculaire et Cellulaire (IGBMC), CNRS (UMR 7104), Inserm U964, Université de Strasbourg, Illkirch Graffenstaden, France; Department of Pathology, Brigham and Women’s Hospital, Boston, Massachusetts, USA; Department of Genetics, Harvard Medical School, Boston, Massachusetts, USA; Harvard Stem Cell Institute, Boston, Massachusetts, USA; Disease Biophysics Group, Wyss Institute for Biologically Inspired Engineering, School of Engineering and Applied Sciences, Harvard University, Cambridge, USA; Departments of Chemistry and Chemical Biology and Physics, Harvard University, Cambridge, MA, USA; Anagenesis Biotechnologies, Parc d’innovation, Illkirch Graffenstaden, France; Aix-Marseille University, CNRS, Institut de Biologie du Développement de Marseille (UMR 7288), IBDM, Marseille; Department of Molecular Biology, University of Texas Southwestern Medical Center, Dallas, TX 75390, USA

**Keywords:** pluripotent stem cells, myogenesis, skeletal muscle, satellite cell, bioengineering, directed differentiation, embryonic stem cell, Duchenne Muscular Dystrophy, dystrophin

## Abstract

Duchenne Muscular Dystrophy (DMD) is a devastating genetic disease leading to degeneration of skeletal muscles and premature death. How dystrophin absence leads to muscle wasting remains unclear. Here, we describe an optimized protocol to differentiate human induced Pluripotent Stem Cells (iPSC) to a late myogenic stage. This allows to recapitulate classical DMD phenotypes (mislocalization of proteins of the Dystrophin-glycoprotein associated complex (DGC), increased fusion, myofiber branching, force contraction defects and calcium hyperactivation) in isogenic DMD-mutant iPSC lines *in vitro*. Treatment of the myogenic cultures with prednisolone (the standard of care for DMD) can dramatically rescue force contraction, fusion and branching defects in DMD iPSC lines. This argues that prednisolone acts directly on myofibers, challenging the largely prevalent view that its beneficial effects are due to anti-inflammatory properties. Our work introduces a new human *in vitro* model to study the onset of DMD pathology and test novel therapeutic approaches.

---

Duchenne Muscular Dystrophy (DMD) is an X-linked muscular dystrophy (affecting 1 in 5000 boys) caused by mutations in the dystrophin gene (aka DMD)(Rahimov and Kunkel, 2013). There is currently no cure for the disease and the only available treatment are glucocorticoids which can prolong the ambulatory phase (Matthews et al., 2016). The dystrophin protein plays a key role in organizing a molecular complex (dystrophin associated glycoprotein complex (DGC)) spanning the sarcolemma at the level of costameres and linking the actin cytoskeleton to laminin and extracellular matrix. In DMD patients, fibers are more sensitive to mechanical stress and experience formation of membrane tears upon muscle contraction (Petrof et al., 1993). DMD mutant myofibers exhibit abnormal calcium homeostasis, displaying higher resting calcium levels (Burr and Molkentin, 2015). The DGC also acts as an important scaffold necessary for the function of several signaling proteins such as the Nitric Oxide Synthase (nNOS) (Brenman et al., 1995). In the early stages of the disease, the degeneration of muscle fibers stimulates regeneration of new fibers from satellite cells, a physiological response that counterbalances fiber loss and maintains a normal muscle function. This increased generation of fibers is accompanied by structural defects such as branching of the newly generated fibers possibly resulting from fusion defects of the regenerating cells (Chan and Head, 2011; Schmalbruch, 1984). As the disease progresses, satellite cell regeneration capacity decreases leading to tissue fibrosis. This myofiber degeneration and fibrosis are considered to be largely responsible for the decrease in muscle strength observed in patients. There is also evidence suggesting intrinsic contractile dysfunction in zebrafish, mouse or dog lacking dystrophin (Lowe et al., 2006; Widrick et al., 2016; Yang et al., 2012). Due to the difficult access to patient muscle fibers, evidence for such contraction defects and their cause and significance for the disease in humans has remained very limited (Fink et al., 1990).

Much of the research on the etiology of DMD as well as preclinical tests for the validation of DMD therapeutic strategies have been carried out in the *mdx* mouse, a spontaneous dystrophin mutant (Partridge, 2013). In the *mdx* mouse myofibers, defects such as branching, and misalignment are detected as early as E13.5 at the beginning of the fetal period (Merrick et al., 2009). A significant limitation of the *mdx* model is that the dystrophy is much less severe and only partly phenocopies the human disease (Partridge, 2013). There is therefore a critical need for a better preclinical model in which the disease can be recapitulated with human myofibers. The recent development of protocols to differentiate human pluripotent stem cells such as iPSCs to skeletal myofibers *in vitro* (Chal and Pourquie, 2017) now offers the possibility to generate DMD models better reflecting the physiology of human cells. Several studies describing the establishment of DMD patient iPSC lines and their differentiation to skeletal muscles have been reported (Piga et al., 2019). However, only a limited set of phenotypes have been analyzed and the impact on skeletal muscle contractility has not been investigated.

Here we describe an optimized myogenic differentiation protocol resulting in significantly improved myofiber maturation from human pluripotent cells *in vitro,* as shown by the expression of all fast myosin isoforms. Using this optimized protocol, we show that muscle fibers derived from two human isogenic iPSC cell lines carrying different DMD mutations engineered in a healthy iPSC line, recapitulate most hallmarks of the DMD phenotype compared to the parental line. These include mislocalization of DGC proteins such as nNOS, branching/fusion defects, and calcium signaling hyperactivation. We also demonstrate that skeletal myofibers derived from the DMD mutant lines exhibit defective contractions, strongly supporting the notion that an intrinsic contractility defect also contributes to the muscle weakness phenotype in DMD patients. Remarkably, this contraction defect can be largely rescued by prednisolone treatment indicating that the drug directly acts on mutant fibers. Finally, these DMD phenotypes are also observed in an iPSC line derived from a DMD patient differentiated using the optimized protocol and they are rescued when restoring the *DMD* coding frame using CRISPR-Cas9, demonstrating their specificity. Thus, our work provides a novel *in vitro* platform to study the etiology of DMD in human myofibers. Our human DMD *in vitro* model will allow for exploration of the early contraction and branching defects caused by absence of Dystrophin at the origin of the pathology and offers a platform for preclinical testing of candidate therapies for this devastating disease.

## RESULTS

### Optimization of the maturation of human iPSC-derived muscle fibers

We have developed a 2-step muscle differentiation protocol for human pluripotent stem cells which first recapitulates the early stages of paraxial mesoderm differentiation followed by myogenesis *in vitro* (Figure 1a) (Chal et al., 2016; Diaz-Cuadros et al., 2020). During the first step (primary differentiation), cells are cultured for 3-4 weeks in a series of different media, resulting in the formation of long striated myofibers interspersed with PAX7-positive myogenic precursors (Chal et al., 2016). To monitor progress through myogenic differentiation in these conditions, we performed RNA sequencing (RNA-seq) of the cultures at day 0, day 8, day 16, day 24 and day 32 of differentiation (Figure 1b). We observed a sequence of expression of myogenic markers starting with *PAX3* at day 8 followed by *PAX7* at day 16. At day 16, we first detected *MYF5, MYOG* and *MYOD1* which peaked later between day 24 and 32 when the marker for fetal (secondary) myogenesis *NFIX* was strongly expressed (Messina et al., 2010). mRNAs for sarcomeric proteins such as α-actinin *(ACTN2)* or titin (*TTN*) were first detected at day 16 and peaked at day 32. At day 32, we also detected expression of both embryonic *(CHRND)* and adult *(CHRNE)* subunits of the acetylcholine receptor. Overall, these data suggest that the primary differentiation protocol recapitulates embryonic and fetal myogenesis *in vitro.* In the second step, which can start after ~20-30 days *in vitro,* the myogenic cultures are dissociated and re-plated in proliferation medium (SKGM), resulting in enrichment in myogenic precursors (Figure 1a) (Chal et al., 2016). After 1-2 days, when the re-plated cells reach confluence, they can be switched to a basal differentiation medium (which contains the Wnt agonist CHIR and KSR (Knockout Serum Replacement)). This results in cultures highly enriched for long striated myofibers after 7 days (Figure 1c). We next analyzed the effect of supplementing the basal medium of these cultures with a TGFβ inhibitor (SB-431542) previously reported to enhance fusion efficiency of muscle fibers (Hicks et al., 2018; Sieiro et al., 2019). When differentiated for 1 week in <u>K</u>SR/<u>C</u>HIR (KC) or in <u>K</u>SR/<u>C</u>HIR/ TGFβ <u>inhibitor</u> (KCTi), the replated cells elongated and rapidly acquired a myogenic phenotype, developing into multinucleated muscle fibers (Figure 1c-d). Fibers generated in KCTi medium appeared thicker with better organized sarcomeres compared to the KC medium (Figure 1c-d). During human fetal development, a burst of glucocorticoid signaling is observed around 7 to 14 weeks, when fetal myogenesis is ongoing (Busada and Cidlowski, 2017). To recreate conditions similar to that experienced by the developing human fetus, we treated the re-plated cultures differentiating in KCTi with prednisolone (<u>K</u>SR/<u>C</u>HIR/<u>T</u>GFb<u>i</u>/<u>P</u>rednisolone (KCTiP)), a synthetic gluco-corticoid hormone previously shown to promote myogenic differentiation in wild type (WT) and *mdx* primary myoblast cultures (Braun et al., 1989). Even though expected myogenic markers were expressed in cultures differentiated in KC and KCTi, we noted a very significant improvement of the morphology of cultures treated with KCTiP with more organized myofibrils (Figure 1c-f, supplementary Figure 1).

**Figure 1:**
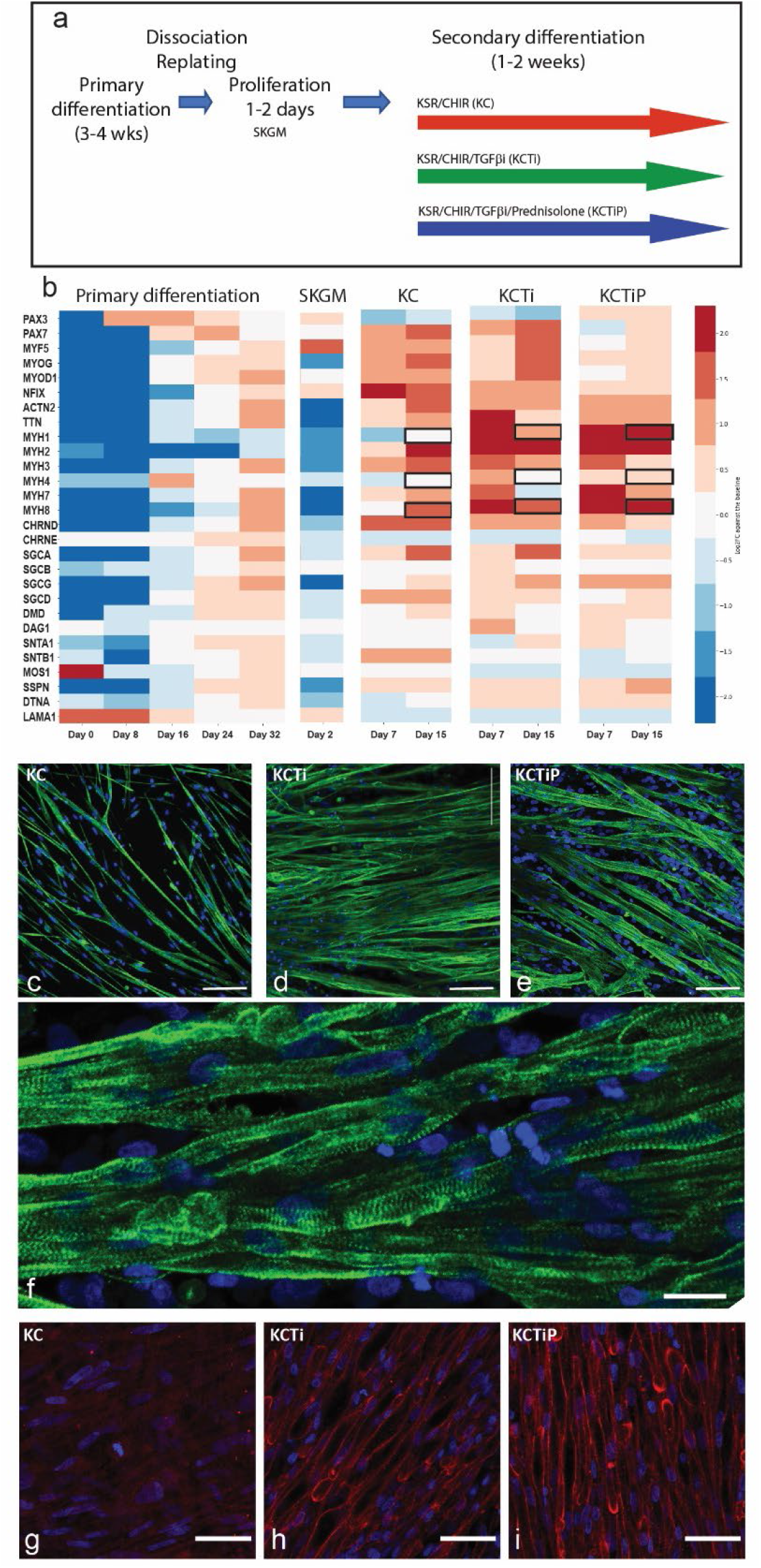
Generation and maturation of iPSC-derived myofibers. (a) Schematic description of the two-step Myogenic differentiation protocol. (b) RNAseq analysis of the myogenic differentiation of wild type human iPSCs *in vitro.* Heat map showing expression levels of selected myogenic markers at different time points during primary differentiation, proliferation in SKGM after replating, and secondary differentiation in KC, KCTi or KCTiP media. (c-f) Titin antibody staining of secondary cultures differentiated for 1 week in KC (c), KCTi (d) and KCTiP (e-f) media. (c-e) Bar: 100 μm. (f) Bar: 25 μm. (g-i) Dystrophin immunocytochemistry analysis following secondary differentiation for 1 week in KC (g), KCTi (h) and KCTiP (i) media. Bar: 50 μm.

We next examined the expression of Myosin Heavy Chain isoforms mRNAs which are sequentially activated during skeletal muscle development (Schiaffino et al., 2015). Embryonic Myosin Heavy Chain *(MYH3)* was strongly expressed at day 16 while a weak expression of Neonatal myosin *(MYH8)* and slow myosin *(MYH7)* was observed at this stage (Figure 1b). Expression of *MYH8* and *MYH7* strongly increased at day 32 in the cultures. During primary differentiation, we only detected low levels of expression of the fast myosin IIa (*MYH1*), IIx *(MYH2)* and IIb *(MYH4)* which are respectively first expressed during fetal, late fetal, and early post-natal stages (Schiaffino et al., 2015). We also performed RNA-seq analysis of the secondary cultures differentiated following re-plating in KC and KCTi for 1 week and compared them to the non-differentiated myogenic progenitors grown in SKGM and to primary differentiation (Figure 1 a-b). When compared to primary differentiation and to KC medium, KCTi induced a higher level of expression of the fast myosins IIa (MYH1), IIx (MYH2) and IIb (MYH4). RNA-seq analysis of cultures in KCTiP show a further increased expression of MYH1, MYH2 and MYH4 compared to cultures treated with KCTi only (Figure 1b). Thus, exposing differentiating human iPSC cultures to glucocorticoids promotes the maturation of skeletal myofibers.

### Generation and differentiation of isogenic DMD mutant iPSC cell lines

We used CRISPR-Cas9 mediated gene editing to establish isogenic cell lines in which DMD mutations were engineered into the wild-type human iPSC line NCRM1. We engineered a full deletion of exon 52 (hereafter named DMDI) and a point mutation introducing a stop codon in exon 52 (named DMDII). We could not detect any Dystrophin protein by western blot and immunohistochemistry in myofibers derived from the two mutant lines compared to the parental line (Figure 2a-d). We did not observe striking differences when comparing the expression of the myogenin, desmin, titin, and α-actinin proteins between skeletal muscle fibers formed from the parental WT cells and the two DMD mutant lines in re-plated cultures differentiated for a week in KCTiP medium (Figure 2e-h).

**Figure 2:**
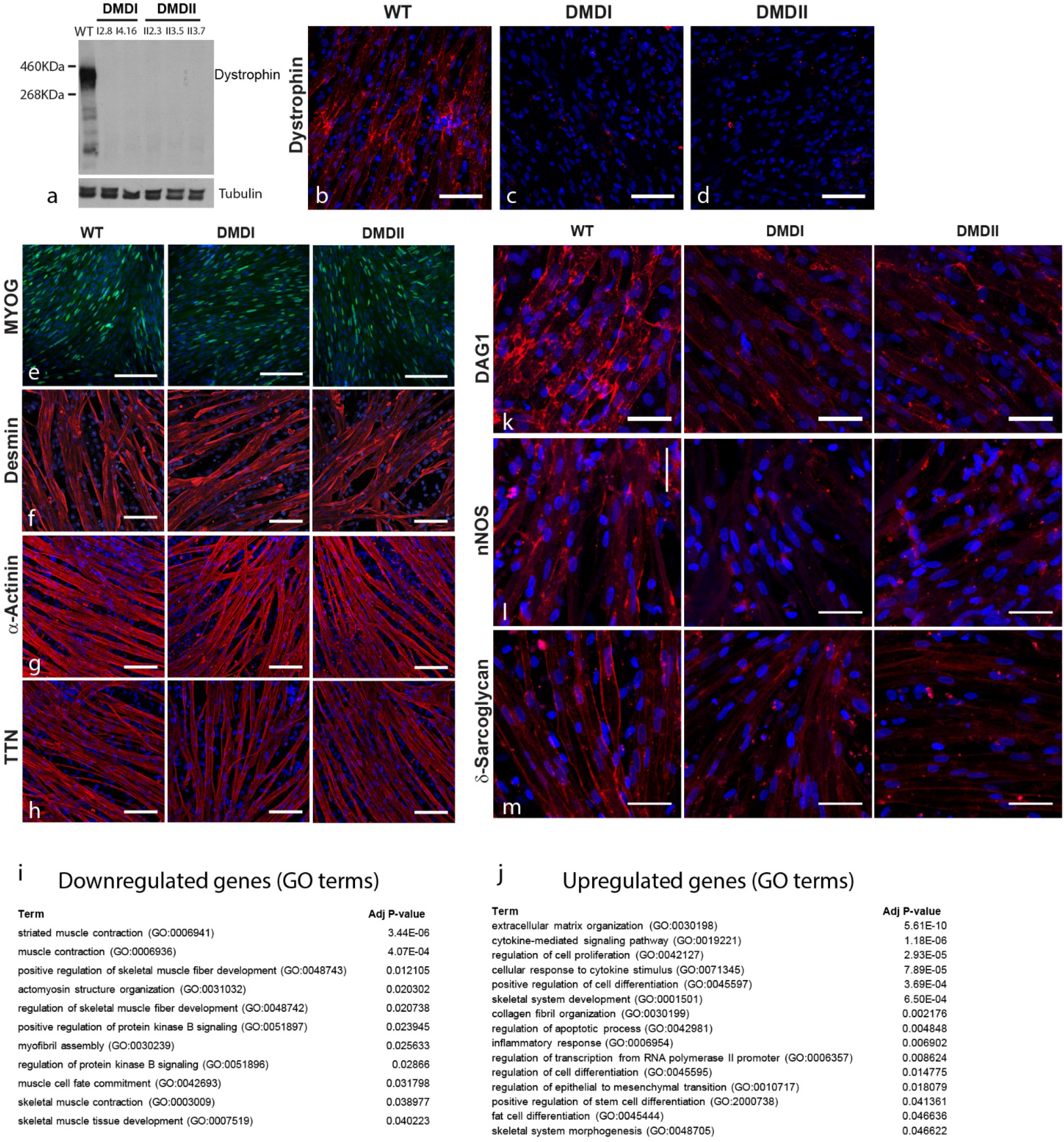
Characterization of isogenic DMD mutant human iPSC lines. (a) Western blot analysis with an anti-Dystrophin antibody in three independent clones of the exon 52 deletion mutant (DMDI) and the exon 52 point mutation mutant (DMDII). Bottom: Tubulin loading control. (b-d) Dystrophin expression detected by immunocytochemistry in secondary cultures differentiated for 1 week in KCTiP medium of WT (b), DMDI (c) and DMDII (d) mutant lines. Bar: 100 μm. (e) Immunocytochemistry analysis of myogenic precursors expressing Myogenin (MYOG) after 2 days of secondary differentiation in SKGM medium of WT (left), DMDI (middle) and DMDII (right) mutant lines. Bar: 200 μm (f-h) Immunocytochemistry of secondary cultures differentiated for 1 week in KCTiP medium of WT (Left), DMDI (Middle) and DMDII (Right) mutant lines. (f) Desmin, (g) α-actinin, (h) Titin (TTN). Bar: 100 μm. (i-j) Significantly enriched GO terms from ‘Biological Process’ in downregulated (i) and upregulated (j) genes in secondary cultures differentiated for 1 week in KC medium of the DMDI and DMDII mutant lines compared to the parental WT iPSCs. (k-m) Immunocytochemistry of secondary cultures differentiated for 1 week in KCTiP medium of WT (Left), DMDI (Middle) and DMDII (Right) mutant lines. (k) DAG1, (l) nNOS, (m) Delta-sarcoglycan. Bar: 50 μm Nuclei (blue) are stained with DAPI. WT: Wild Type

Next, we used RNA-seq to compare the transcriptome of myogenic cultures of the two mutant lines with the parental line after SKGM amplification and differentiation for 1 week in KC medium. 532 genes were downregulated while 723 genes were upregulated in both DMD lines with a fold change >2 and an FDR< 0.01, when compared to the WT cells (Supplementary Table 1). Gene Ontology (GO) analysis of the differentially expressed genes revealed that down-regulated genes were primarily enriched in GO terms related to muscle including “striated muscle contraction” or “positive regulation of skeletal muscle development” as well as “regulation of protein kinase B signaling” (Supplementary Table 2 Figure 2i). Transcription factors found in these categories included MYOD1, MYF6, MYOG and MEF2C (Supplementary Table 3). GO terms primarily enriched in the upregulated genes in DMD cultures included “extracellular matrix organization”, “cytokine-mediated signaling pathways” and “fat cell differentiation”, which are in line with the inflammation and fibrosis and fat cell infiltration detected in patients (Supplementary Table 2, Figure 2j).

### Mislocalization of proteins of the Dystrophin-associated Glycoprotein Complex in DMD mutant iPSC lines

When parental WT cells were differentiated in KCTiP medium, muscle fibers strongly expressed Dystrophin compared to cells cultured in KC or KCTi conditions (Supplementary Figure 2). Moreover, stronger expression of the components of the DGC complex dystroglycan (DAG1) and gamma-sarcoglycan was also detected when prednisolone was present in the differentiation medium (Supplementary Figure 2). In both DMD mutant lines, the DGC proteins nNOS, DAG1 and gamma-Sarcoglycan were largely absent from the sarcolemma of the myofibers, where they are localized in the parental WT line (Figure 2 k-m). Overall, proteins of the DGC were mis localized and downregulated in DMD mutant fibers whereas other membrane associated proteins such as NCAM1 were not affected (Supplementary Figure 3). Therefore, misexpression of DGC proteins reminiscent of the phenotype of DMD patients (Brenman et al., 1995; Janghra et al., 2016) is observed in DMD skeletal myofibers differentiated *in vitro* in KCTiP medium.

### Increased branching and fusion of differentiated Dystrophin-deficient fibers

To test whether human DMD fibers generated *in vitro* exhibit a branching phenotype, myogenic cultures from the two DMD isogenic lines and from the WT parental line were dissociated at 3-weeks and re-plated. After 24h, progenitors were transfected at low efficiency with membrane GFP and nuclear mCherry constructs. This allowed us to permanently mark isolated myofibers within the population, and then to quantify the number of branching points as well as nuclei within individual fibers. After 1 week of differentiation in KC medium, fibers with no branches were observed in 77% of cases, with an average of 0.35 branching points per fiber for the entire WT population (Figure 3a, j-k). In contrast, in DMD-derived fibers, an average of 0.59 branching points per fiber were observed for both mutant lines, while around 65% and 62% of fibers remained unbranched for DMDI and DMDII respectively (Figure 3a-c, j-k). When fibers were differentiated in the presence of TGFβ inhibitor (KCTi), a markedly significant increase in branching points was observed when compared to KC medium (Figure 3d-f, j-k). WT fibers differentiated in KCTi contained an average of 0.94 branching points, with more than half of the fibers (53.2%) being bifurcated at least once. Dystrophin-deficient fibers averaged a significantly higher number of branching points when compared to wild-type cells (1.42 and 1.25 for DMDI and DMDII respectively) in KCTi. We then investigated the effect of prednisolone on myofiber branching. WT fibers grown in KCTiP medium contained no branching points in 79% of cases, a proportion higher than those of WT fibers grown in KCTi medium (Figure 3g-k). Interestingly, Dystrophin-deficient fibers differentiated in KCTiP also showed fewer branching points than those grown in KCTi media although they maintained a significantly higher number of branches when compared to WT (Compare Figure 3d-f and g-i).

**Figure 3:**
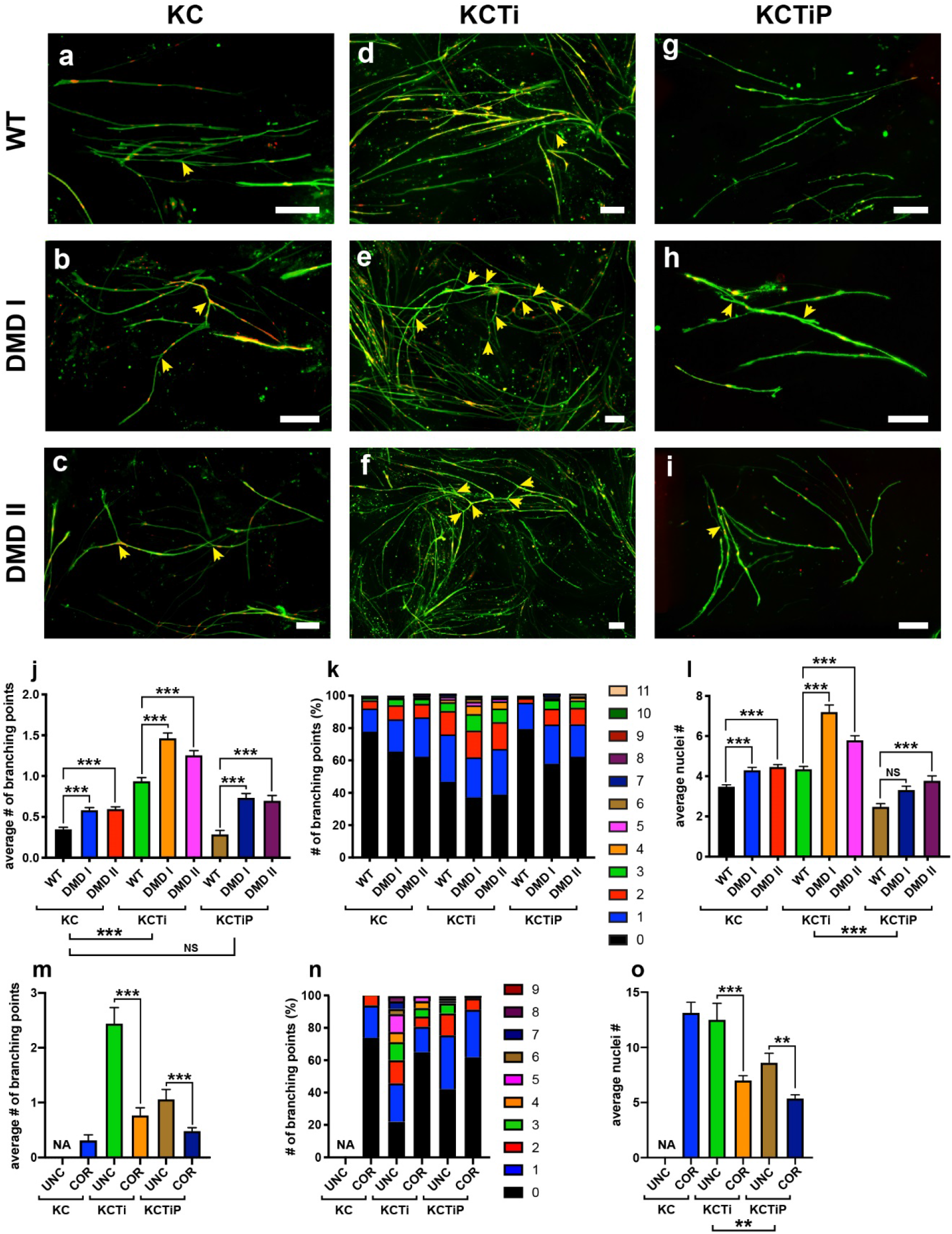
Myofibers differentiated *in vitro* from Dystrophin-deficient iPSC lines exhibit increased branching defects. (a-i) Isolated fibers from secondary cultures WT (a, d g) and DMDI (b, e, h) and DMDII (c, f, i) iPSC lines labeled with membrane GFP (green) and mCherry (red) signals differentiated for 1 week in KC (a-c), KCTi (d-f) or KCTiP (g-i) medium. Yellow arrowheads indicate branching points. Bar: 20μm (j-k) Quantitative analysis of the number (#) of branching points in WT and DMD isogenic lines in the different culture media. ***: p<0.0001, NS: p>0.05. Bars show mean +/− SEM. (l) Quantitative analysis of the number of nuclei per fiber in WT and DMD isogenic lines in the different culture media. ***: p<0.0001, NS: p>0.05. Bars show mean +/− SEM. (m-n) Quantitative analysis of the number of branching points in TX1-UNC and TX1-COR isogenic lines in the different culture media. ***: p<0.0001. Bars show mean +/− SEM. No data is shown for TX1-UNC in KC (NA) as myogenic differentiation from these lines was poorly efficient in this condition. (o) Quantitative analysis of the number of nuclei per fiber in TX1-UNC and TX1-COR isogenic lines in the different culture media. ***: p<0.0001. Bars show mean +/− SEM. Kruskal-Wallis non-parametric ANOVA test with planned multiple comparisons. WT: Wild Type. SEM: Standard error of the mean.

We also investigated how the absence of Dystrophin impacts the fusion of myofibers. Nuclei labeled by mCherry were counted in isolated fibers in cultures of WT and DMD mutant cells differentiated in KC, KCTi or KCTiP. In all three different conditions, we observed a significant increase in the number of nuclei in the DMD mutants compared to WT fibers (Figure 3l). To confirm the specificity of the branching and fusion defects in a different genetic background, we used a human patient-derived iPSC line harboring an intronic point mutation in intron 47 of the DMD gene (TX1-Unc) and an isogenic line in which the Dystrophin coding frame was restored by CRISPR-Cas9 editing (TX1-Cor) (Long et al., 2018). Both lines could differentiate efficiently into myofibers expressing sarcomeric proteins in KCTi and KCTiP (Supplementary Figure 4). No expression of Dystrophin and downregulation of DAG1 was observed in the TX1-Unc line while expression of these proteins in the TX1-Cor line was similar to WT (Supplementary Figure 4). The number of branching points and the number of nuclei per fiber was significantly higher in the uncorrected TX1-Unc line in KCTi than in the corrected line (Figure 3m-o). Remarkably, the number of branching points and of nuclei per fiber could be reduced in both lines by prednisolone treatment (Figure 3m-n). Strikingly, prednisolone treatment rescued the branching phenotype in the patient iPSC line to the level of the corrected line treated or not with prednisolone (Figure 3m). Thus, our data show that absence of Dystrophin leads to an increase in myofiber branching and fusion in three different DMD mutant isogenic lines in various differentiation conditions. Remarkably, excessive branching and fusion can be reduced by treating the differentiating cultures with prednisolone.

### Force contraction defects displayed in Dystrophin-deficient iPSC derived fibers

To measure the impact of loss of Dystrophin on force contraction, we engineered contractile myogenic tissues from the WT and DMD mutant lines (Figure 4a)(Chal et al., 2016; Nesmith et al., 2016). Myogenic cultures of the WT and the DMDI and DMDII mutant iPSC lines were dissociated and seeded onto thin elastomeric gelatin substrates, which were micro-molded with line patterns to promote cell alignment (Supplementary Figure 5). The re-plated cells were first cultured for 1-2 days in SKGM until they reached confluence and then were differentiated for a week in KCTi medium. In these conditions, myocytes self-organized into continuous multi-nucleated myofibers, forming muscular thin films (MTFs) with an average myofiber thickness of ~15 μm (Figure 4a-d, Supplemental Figure 6). Immunofluorescence staining of α-actinin revealed highly aligned sarcomeres in both wild-type and DMD mutant muscle fibers (Figure 4b-d).

**Figure 4:**
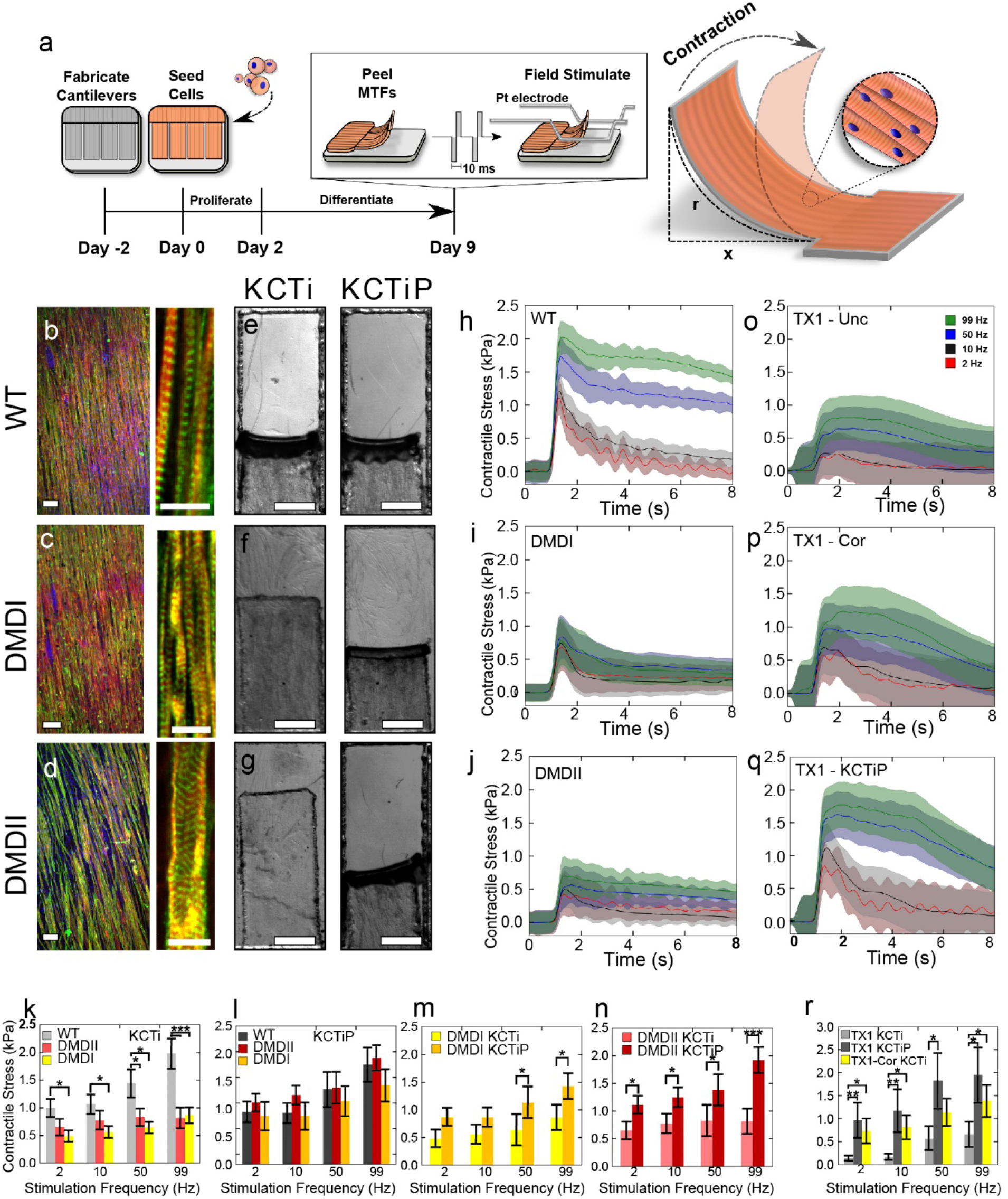
Dystrophin mutant fibers derived *in vitro* exhibit contractile defects. (a) Experimental Protocol and schematic illustration of muscular thin film (MTF) assay, for measuring contractile force. (b-d) Representative immunofluorescent micrograph of skeletal muscles grown on micro-molded gelatin substrates, showing aligned confluent tissues (b-WT, c-DMD I, d–DMD II) (DAPI: Blue, α-actinin: Green, actin: Red), with high magnification inset showing sarcomere expression. (Scale bars: 100 μm left, 10 μm right). (e-g) Bright field micrographs of MTF cantilevers (Top-down view) for WT (e), DMD I (f) and DMD II (g) myofibers cultured in KCTi (left) and KCTiP (right) media after stimulation (99 Hz), showing increased contraction in DMD cells after exposure to prednisolone. (Scale bars: 1 mm). (h-j) Skeletal muscle contractile force as a function of stimulation time for WT (h), DMDI (i), and DMDII (j) mutant iPSC lines, demonstrating a positive force frequency relationship in the absence of dystrophin disruption (cells stimulated between 1.5-8 seconds, at 2-99Hz). (k-n) Comparison of peak contractile stresses generated by MTFs paced at 2-99 Hz. (Scale bars: 1 mm), showing a stronger contractile phenotype for WT cells (k), but a recovery of contractile phenotype in the presence of prednisolone treatment (KCTiP) (l-n). (n> 22, Supplementary Table 3). * = p<0.05, ** = p<0.01, *** = p<0.001). (o-q) Skeletal muscle contractile force as a function of stimulation time for DMD patient-derived cells (uncorrected -o, q, Corrected - p) in KCTi (o-p) and KCTiP (q) media (cells were stimulated between 1.5-8 second. at 2-99Hz). (r) Peak contractile stress generated by patient derived cells paced at 2-99 Hz. (Scale bars: 1 mm), showing a recovery of contractile phenotype for patient cells treated with prednisolone or corrected with CRISPR-Cas9. (n> 15, Supplementary Table 4). Pairwise t-tests. All error bars given as the standard error of the mean.

Muscle constructs were field stimulated using a frequency sweep over 2-99Hz, transitioning between twitch stresses to tetanic contractions. Film deformation was recorded using a stereomicroscope (Figure 4e-g) and changes in tissue radius, *r*, were mapped to contractile stresses. For WT cultures, greater stimulation frequencies lead to increased sustained contractile stresses between 1,000-2,000 Pa (Figure 4h). For DMDI and DMDII, increases in stimulation frequency produced only a mild, non-statistically significant increase in contractile stress between 500-900 Pa (Figure 4i-j). For WT cells, stimulation at 99Hz yielded an approximate specific tensile strength of 180 ±82 kPa, which is on par with human *in vivo* muscle measurements (Maganaris et al., 2001). Thus, the DMD cell lines showed overall lower contractile stresses and a minimal force-frequency response as compared to WT muscle differentiated *in vitro* (Figure 4k).

DMDI and DMDII cultures were next differentiated in the presence of prednisolone in KCTiP medium (Supplementary Figure 7). After one week, DMD-derived muscle fibers showed a restored contractile function, yielding positive force-frequency relationships with contractile stresses between 1,000-2,000 Pa and 850-1,400 Pa for DMDII and DMDI respectively (Figure 4e-g, l-n). These levels were comparable to those of control WT cantilevers, cultured with or without prednisolone (Figure 4l, Supplementary movie 1).

Additionally, we compared force contraction between the TX1-Unc and TX1-Cor isogenic pair. MTFs derived from the TX1-Unc and TX1-Cor lines formed aligned myofibers (Supplemental Figure S7) but displayed distinct contractile phenotypes, with greater contraction stress generated by the corrected line (700-1,400 Pa) compared to the parental line (150-650 Pa) (Figure 4o-r). Furthermore, prednisolone treatment of the parental TX1-Unc line rescued force contraction to the level of the corrected TX1-Cor line (950-1950 Pa) (Figure 4o-q, Supplementary movie 2). These data demonstrate that DMD mutation leads to defects in force contraction in myofibers differentiated *in vitro,* and that these contractile defects can be largely rescued by prednisolone treatment.

### Dystrophin-deficient fibers show Ca^2+^ hyper-excitability

We next combined optogenetics and Ca^2+^ imaging to study the dynamics of Ca^2+^ handling in our DMD mutant myofibers differentiated *in vitro.* We used a lentivirus to infect proliferating myogenic precursors in SKGM (Figure 1a), to express a blue-light sensitive channel-rhodopsin CheRiff (Figure 5a)(Hochbaum et al., 2014). After 7-10 days of differentiation in KCTi medium, we incubated the myofiber cultures with the Ca^2+^-sensitive dye CaSiR-1 AM. We next mapped Ca^2+^ responses across large (4 mm x 4 mm) cultures of myofibers using a custom-built ultra-widefield microscope (Figure 5b-c). In all experiments, we observed a Ca^2+^ ‘hyperexcitability’ phenotype for myofibers lacking Dystrophin, in which the amplitude of Ca^2+^ responses was considerably higher for the mutant fibers than for healthy controls across the range of stimulus frequencies (Figure 5d-f, Supplementary movie 3). For isogenic cultures, we observed statistically significant differences between WT and DMDII samples for all tested stimulus frequencies, and between WT and DMDI samples for stimulus frequencies between 2 and 20 Hz (Figure 5d). Notably, DMD fibers showed slower relaxation kinetics than did healthy fibers. The patient-derived line similarly showed statistically significant differences in Ca^2+^ responses at each frequency tested (Figure 5e, Supplementary movie 4), with greater Ca^2+^ responses in the parental TX1-Unc line compared to the CRISPR-corrected TX1-Cor line. We further validated the patient-derived cell line results by repeating the experiment using a different Ca^2+^-sensitive dye (BioTrack 609), and by normalizing responses relative to an ionomycin treatment (Figure 5f, Supplementary movie 5). This experiment again showed elevated Ca^2+^ responses in Dystrophin-deficient fibers, consistent with the Ca^2+^ hyperexcitability phenotype. Each experiment was consistent with pathophysiological elevated and sensitized Ca^2+^ responses in dystrophic fibers. Notably, observation of elevated “gain” of Ca^2+^ signaling in response to a frequency ramp, as well as differences in relaxation kinetics, suggest an involvement of Ca^2+^ handling feedbacks beyond an increase in leakage Ca^2+^ currents across the plasma membrane. These results demonstrated that iPSC-derived skeletal myofibers can recapitulate phenotypes of Ca^2+^ handling in both isogenic and patient derived cell lines.

**Figure 5:**
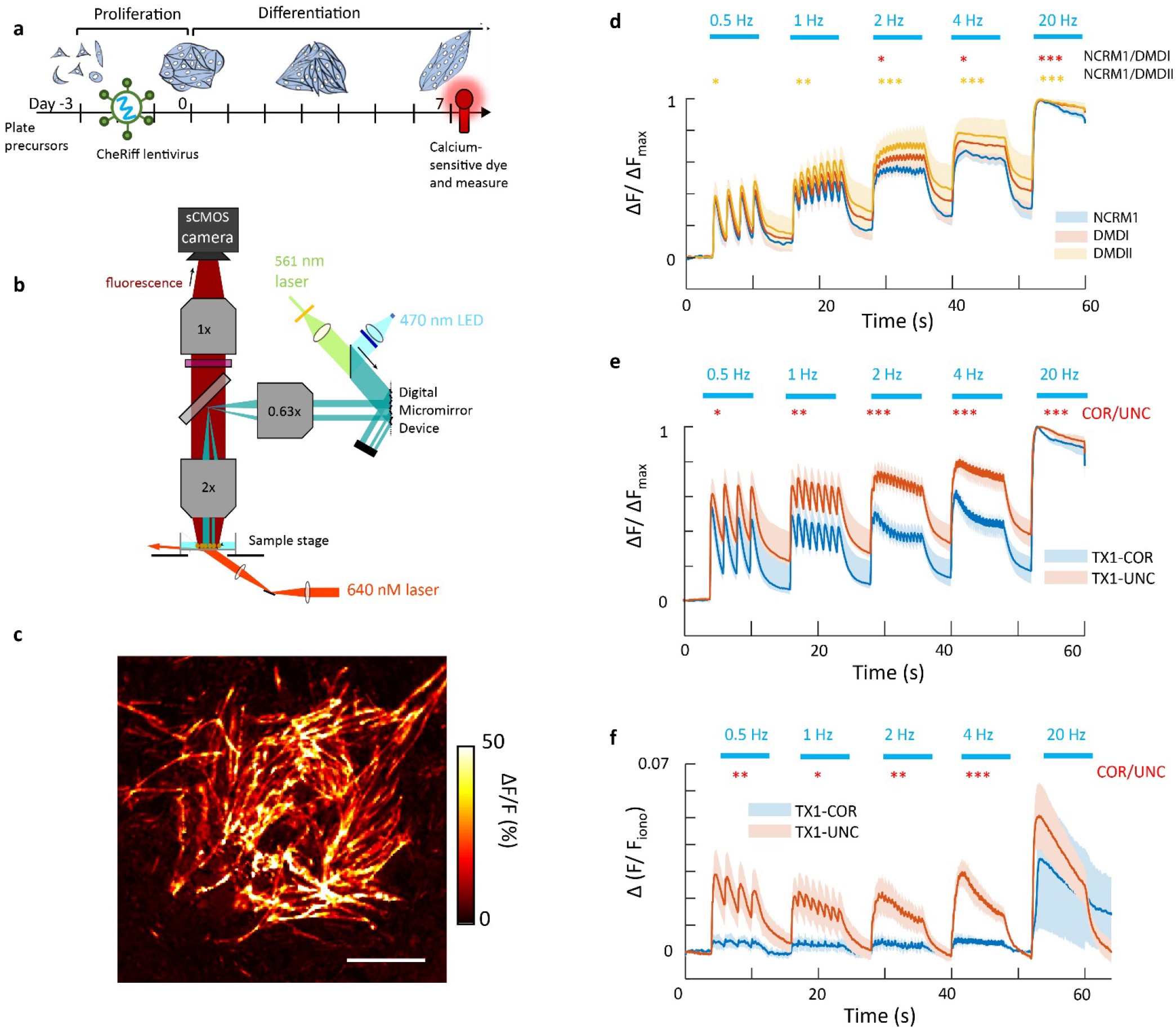
All-optical profiling of Ca^2+^ responses in healthy and dystrophic human iPSC myofibers. (a) Schematic of experimental timeline. Myocyte precursors are plated at low density in growth medium and inoculated with a lentiviral vector encoding CheRiff after 24 h. After 72 h, cells have reached confluence and are switched to differentiation media and cultured for 7 additional days in KCTi medium. After 7 days of differentiation, cultures are incubated with the Ca^2+^ sensitive dye CaSiR-1 AM and measured. (b) Diagram of optical setup. Red Ca^2+^-sensitive dyes are excited using oblique illumination, and Ca^2+^-sensitive fluorescence in the near infrared is collected with a high-NA widefield objective. CheRiff stimulation is spatially targeted using a digital micromirror device. (c) Example image of blue-light induced Ca^2+^ response in a dish of iPSC cell-derived myocytes. Scale bar = 1 mm. (d) Profiling Ca^2+^ response as a function of blue light drive. Simultaneous differentiations of WT, DMDI, and DMDII cell lines (N=6 dishes of each) were characterized via their Ca^2+^ response to optogenetic stimulation across a range of drive frequencies (0.5 Hz to 20 Hz). Average traces reveal statistically significant differences between DMDI and NCRM1 lines (red asterisks) and between DMDII and NRCM1 lines (yellow asterisks). DMDI and DMDII lines showed no significant difference in Ca^2+^ responses. (e) Same experiment as in (d), but with parallel differentiations of a patient derived iPSC line (TX1-UNC) and a corrected comparison (TX1-COR) (N=6 dishes of each). For patient-derived cells, N=6 samples were analyzed for both TX1-COR and TX1-UNC for the first replicate, (f) Replicate experiment comparing TX1-COR and TX1-UNC patient derived cultures, in which Ca^2+^ signals are imaged with BioTracker 609 and normalized relative to responses to ionomycin application (10 μM). N=4 samples were analyzed for each condition in the second replicate. Paired two-sample t-tests. Confidence intervals of p<0.05 (*), p<0.01(**), and p<0.001(***).

## DISCUSSION

Previous *in vitro* models based on myofibers differentiated from DMD patient iPSC lines have led to discordant results (Piga et al., 2019). A limitation of several of these studies is the comparison of iPSC lines from patients to lines from healthy subjects. The inherent variability in the differentiation potential of individual lines (Osafune et al., 2008) is highly problematic as it can confound phenotypical studies. This problem can be circumvented by using isogenic lines in which a disease-causing mutation is introduced in a healthy parental line whose differentiation properties are well characterized. Here, we report for the first time the engineering of human DMD mutant iPSC isogenic lines from a healthy WT line. We generated a deletion of exon 52 and a point mutation in exon 52 using CRISPR-Cas9 editing in the NCRM1 WT iPSC line. Thus, the phenotype of the engineered lines can be directly compared to the WT parental line whose myogenic differentiation has been well characterized (Chal et al., 2016).

A second limitation from previously reported iPSC-based DMD models lies in the immature status of the myofibers generated *in vitro* using current myogenic differentiation protocols. Here, we describe a novel method which significantly increases myofiber maturation over existing protocols. While current strategies can result in differentiation of myogenic cells up to the embryonic to fetal transition (Al Tanoury et al., 2020; Xi et al., 2020), our improved method results in well-organized myofibers with significant activation of the fast myosin IIa (*MYH1*), IIx *(MYH2)* and IIb *(MYH4),* which are respectively first expressed during fetal, late fetal, and early post-natal stages (Schiaffino et al., 2015). While most of DMD pathological landmarks have been defined during post-natal stages, the primary events leading to these defects likely happen during fetal development. The macroscopic architecture of fetal DMD muscles appears similar to normal fetal muscle, but myofiber defects and abnormal Ca^2+^ signaling are already observed in DMD fetuses (Emery, 1977; Emery et al., 1979; Farini et al., 2016). Signs of degeneration and regeneration of the myofibers become conspicuous soon after birth, before clinically detectable symptoms (Pearson, 1962). Studying these early stages of disease development is challenging due to the very limited access to DMD fetuses and thus these early defects are poorly understood. Our system can therefore help understanding the earliest defects resulting from DMD absence in human patients.

In contrast to some studies (Choi et al., 2016; Moretti et al., 2020), we show that in our conditions, the morphology and expression of myogenic markers of differentiating myogenic DMD-mutant iPSC lines is very similar to that of the WT parental NCRM1 line. The most striking morphological phenotype we observed in the differentiating DMD myofibers is abnormal branching, a defect which has been reported in muscle biopsies from boys with DMD and in myofibers of dystrophin-deficient *mdx* mice (Chal et al., 2015; Chan and Head, 2011; Schmalbruch, 1984). The increased branching of DMD mutant myofibers observed *in vitro* was accompanied with an increased number of nuclei per fiber suggesting increased fusion in DMD fibers. This supports the hypothesis that abnormal branching could result from increased fusion required by the sustained regeneration caused by myofibers death in DMD patients (Chan and Head, 2011).

Using a tissue engineering approach in which myofibers are seeded on soft cantilevers (Nesmith et al., 2016), we observed a significant decrease in force contraction in the skeletal myofibers derived from DMD mutant iPSC lines compared to isogenic lines expressing Dystrophin. Such a contraction defect has previously been observed in myofibers derived from patient’s primary myoblasts using the same platform (Nesmith et al., 2016). Such tissue engineered models have been previously used to model cardiac disease including DMD (Long et al., 2018; Wang et al., 2014) and can provide several advantages when compared to *in vivo* models. First, the ability to use isogenic cell lines avoids potential changes in cellular contractility associated with differing genotypes, which can impact baseline muscle growth and tissue development (Costa et al., 2009). Second, this model can be used to directly test human derived cells and subsequent pathophysiology, which have been difficult to recapitulate in animal models (McGreevy et al., 2015). Additionally, tissue engineered models provide the capability of being used for personalized medicine applications, where potential therapeutic interventions are tested against a patient’s own cells. This is especially important in the case of DMD patients, where the dystrophin gene can be disrupted in one of several “hotspot” regions (Esposito et al., 2017), meaning that individual patients may require distinct treatment regimes. Importantly our observations demonstrate that human DMD mutant myofibers exhibit an intrinsic defective contraction defect as observed in zebrafish, mouse or dog lacking dystrophin (Lowe et al., 2006; Widrick et al., 2016; Yang et al., 2012). Our *in vitro* system offers a unique opportunity to understand the cause of this defect and to search for therapeutic strategies to correct it.

The molecular mechanism through which dystrophin loss of function affects Ca^2+^ signaling remains controversial. Here we apply for the first time optogenetics to study Ca^2+^ signaling in human DMD myofibers. We show that *in vitro* differentiation of skeletal myofibers can efficiently recapitulate the Ca^2+^ hyperexcitability phenotype of Dystrophin-deficient fibers. Our data are consistent with the Ca^2+^ handling defects observed in differentiated fibers obtained by forced expression of MyoD in DMD Patient iPSC *in vitro* (Shoji et al., 2015). The hyperexcitability phenotype could result from disruption of the dystrophin-glycoprotein complex leading to increased Ca^2+^ leakage currents (Fong et al., 1990; Franco and Lansman, 1990). It is also possible that feedbacks involved in excitation-contraction coupling and Ca^2+^-induced Ca^2+^ release are dysregulated in DMD myofibers. For example, the sarco/endoplasmic reticulum Ca^2+^ ATPase (SERCA) has been reported to be dysregulated in DMD models, leading to a lack of Ca^2+^ removal from the cytosol (Voit et al., 2017). This would be consistent with our observation of slower Ca^2+^ relaxation kinetics after excitation in DMD lines. Thus, our work introduces a powerful new system to study Ca^2+^ handling in dystrophic myofibers.

Glucocorticoids are part of the standard of care for DMD patients in which they increase force and prolong ambulation (Matthews et al., 2016). The mechanism of action of glucocorticoids underlying their beneficial effect in patients has not been elucidated yet. The positive effect of glucocorticoids on patients is often attributed in part to their immunosuppressive properties (Kissel et al., 1991; Wehling-Henricks et al., 2004). One expected consequence of treatment is a decrease of inflammation associated to degeneration, leading to a slowing down of fibrosis progression and an improvement of muscle function. The glucocorticoid effects are paradoxical because these steroids can also trigger muscle atrophy (Kanda et al., 2001). Prednisolone can also improve myofiber maturation in primary myotubes cultured *in vitro* suggesting that glucocorticoids might also act directly on muscle function (Braun et al., 1989; Sklar and Brown, 1991). In the *mdx* mouse and in patients, glucocorticoid treatment leads to metabolism reprogramming associated to improved performance of muscles (Quattrocelli et al., 2019). Here we demonstrate that Prednisolone can rescue the branching, fusion and force contraction phenotypes in three different DMD mutant iPSC lines *in vitro.* Remarkably, an increase in force contraction is not observed when the WT parental lines are treated with Prednisolone. Thus, our data suggest that prednisolone acts directly on the Dystrophin-deficient myofibers to improve the pathological phenotype. Use of glucocorticoids is problematic in patients as it triggers undesirable side effects such as obesity or mood disorders (Matthews et al., 2016). However, moving forward our *in vitro* system provides an ideal platform to dissect the molecular action of glucocorticoids on myofibers. This will eventually make possible the search for alternative therapies preserving the beneficial effect of glucocorticoids, without the side effects.

## Supporting information

Supplementary Move 2

## Acknowledgements

We thank Lou Kunkel, Emanuela Gussoni and Felipe Leite for critical reading of the manuscript and Suvi Aivio for technical help in the early stages of the project. This work was supported by a strategic grant from the French Muscular Dystrophy Association (AFM) to O.P and by a post-doctoral grant to D.M. Additionally, this work was sponsored in part by the John A. Paulson School of Engineering and Applied Sciences at Harvard University, the Wyss Institute for Biologically Inspired Engineering at Harvard University, and Harvard Materials Research Science and Engineering Center grant DMR-1420570 and an ODET fellowship to J.Z.. HMM and AEC were supported by the Howard Hughes Medical Institute. F.M. is supported by ANR grant ANR-18-CE45-0016 (to B.H.H.).

## Author contributions

Z.A.T. designed and performed differentiation experiments, analyzed data and coordinated the project with O.P. J.Z performed the force contraction measurements. J.R performed differentiations, RNA sequencing and immunostaining experiments. D.S-M. quantified the branching phenotype. H.M. performed the analysis of calcium regulation. T.C. generated the isogenic CRISPR/Cas9-mutant lines in NCRM1 with help of F.B. and C.F. A.H. identified the prednisolone effect on maturation. FM and B.H.H analyzed the RNA-seq data, JC pioneered the force contraction experiments with A.P.N., S.G. and E.W. helped with differentiation experiments. R. B-D and E.O. provided the DMD patient iPSC line and its CRISPR-corrected version. A.C. supervised the calcium experiments. K.K.P. supervised the force contraction experiments. O.P. supervised the project and wrote the manuscript.

## MATERIALS AND METHODS

### iPSC cell maintenance and differentiation

#### Maintenance

Human iPSC cells were cultured as described previously (Chal et al., 2016; Chal et al., 2015). Briefly, cells were cultured on Matrigel (BD Biosciences)-coated dishes in mTesR1 media (Stem Cell Technologies). Cells were passaged as aggregates or as single cells. The NCRM1 human iPSC line (RUCDR, Rutgers University) and its engineered derivatives were tested mycoplasma-free.

#### Differentiation

Serum-free myogenic primary differentiation of human iPSC clones was performed as described previously (Chal et al., 2016; Chal et al., 2015b). For secondary differentiation purposes, 3-week old primary myogenic cultures generated from iPSCs were dissociated as described and myogenic progenitors were replated at a density of 35-40,000/cm^2^ onto Matrigel (Corning, Cat#354277)-coated dishes in skeletal muscle growth media (SKGM-2, Lonza CC-3245) with 10 μM ROCK inhibitor (#1254, R&D Systems) (Chal, Al Tanoury et al 2016). After 24 hours, medium was changed to SKGM-2 media without ROCK inhibitor. Cultures were allowed to proliferate for 1-2 days, at which point they reached ~90% confluence. Cultures were then induced for myogenic differentiation with DMEM/F12 supplemented with 2% knock-out serum replacement (Invitrogen, Cat. # 10828028), 1 μM Chiron (Tocris, Cat. # 4423), 0.2% Pen/Strep (Life Technologies, Cat. # 15140122), 1x ITS (Life Technologies, Cat. # 41400045), with or without 10 μM of the TGFβ inhibitor SB431542 (Tocris, Cat#1614) (KCTi) or 10 μM of Prednisolone (Sigma Aldrich, cat. # P6004) (KCTiP). Following induction, differentiation medium was changed on days 1 and 2 and then was refreshed every other day for 1 week.

### Generation of isogenic DMD mutant cell lines

#### sgRNA design and Cas9 vector assembly

To generate DMD cell line lacking exon 52, NCRM1 cells were transfected using two pSpCas9 (BB)-2A-GFP plasmids: one containing a guide targeting the 5’ intron flanking the exon 52 of the DMD gene and one containing a guide targeting the 3’ intron flanking the exon 52 (Supplementary Table 5). Transfected GFP-positive single cells were sorted by flow cytometry and seeded at very low density in conditioned media. Clone screening was performed by PCR using primers flanking the deleted region (see Supplementary Table 5). Clones exhibiting a perfect repair by non-homologous end joining were selected after sequencing and named DMDI.

To generate the DMD cell line exhibiting a stop codon within exon 52, NCRM1 cells were transfected using a pSpCas9 (BB)-2A-GFP containing a guide targeting exon 52 and a ssODN (Integrated DNA Technologies) containing the mutated region (Supplementary Table 5). Genetic editing consists here in replacing the original DNA sequence TTG GAA GAA CTC ATT ACC by the mutated sequence CTA GAG <u>GAG CTC</u> ATA **TGA**, containing silent mutations creating a SacI restriction site (underlined) and a stop codon (in bold). Transfected GFP-positive single cells were sorted by flow cytometry and seeded at very low density in conditioned media. Positive clones were screened by PCR using the SacI restriction enzyme and, after sequencing, clones exhibiting perfect homology directed repair were selected and named DMDII.

Cas9 target sites were identified using the online CRISPR design tool (http://tools.genome-engineering.org). Briefly, DNA sequences flanking exon 52 of the human DMD gene were used for designing the sgRNAs. Several pairs of sgRNAs targeting either the top or the bottom strands of genomic DNA were selected and tested. For each target, a specific Cas9 vector was made. Briefly, the Cas9 vector pSpCas9 (BB)-2A-GFP (pX458; Addgene plasmid ID: 48138) was digested using BbsI and a pair of phosphorylated and annealed oligos (20 bp target sequences) were cloned into the guide RNA locus as described (Ran et al., 2013). All vectors were sequenced to ensure the presence of the right sequence.

#### Nucleofection

cells were dissociated using Accutase (Stemcell Technologies) and 8 × 10^5^ cells were electroporated using 5 mg of total DNA (Ratio 1:1) and the Amaxa Nucleofector kit (Lonza) as described (Ran et al., 2013). 24h post transfection, cells were dissociated and sorted for the expression of Cas9-GFP by fluorescence-activated cell sorting and expanded clonally at low densities. Later, clones were picked up for PCR screening and expansion.

### Bulk RNA-seq analysis

#### Sample collection

NCRM1 line was differentiated into myogenic cultures as described in (Chal et al., 2016) and cells were harvested on day 0, day 8, day 16, day 24 and day 32 of differentiation. For secondary differentiation, primary cultures were dissociated at day 21 and replated as described above in SKGM. Samples were collected after two days in SKGM (SKGM-2, Lonza CC-3245) culture. The cultures were further differentiated in three different conditions - KC, KCTi and KCTiP. For each condition, cells were harvested after 7 and 15 days of differentiation. For each time point samples were collected from 3 independent experiments. RNA was isolated using NucleoSpin® RNA kit (740955, Macherey and Nagel) following manufacturer’s protocol. RNA libraries were prepared using Roche Kapa mRNA Hyper Prep and sequencing was performed on Illumina NextSeq 500 Sequencing platform.

Data from isogenic DMD versus parental iPSC lines, and from the wild-type iPSC differentiation assays were analyzed independently. For both datasets, we used STAR (v2.5.1b) (Dobin et al., 2013) to map sequenced reads to the genome; gene counts were quantified using featureCounts (1.6.2) (Liao et al., 2014). Data from isogenic DMD and parental iPSC lines were mapped to the reference genome assembly UCSC hg19; for differential expression analysis of isogenic DMD iPSC cell line vs WT parental cell lines, we used DESeq2 (v 1.22.2) (Love et al., 2014). Genes were defined differentially expressed when the fold-change (FC) absolute value was greater than 2 and the false discovery rate lower than 0.01. Gene Ontology (GO) enrichment analysis was performed on the differentially expressed genes using EnrichR (Kuleshov et al., 2016).

Sequenced reads coming from the myogenic differentiation protocol were mapped to the reference genome assembly GRCh38 release 77 from ENSEMBL. To produce heat maps for genes of interest from the muscle differentiation assays, we first normalized read counts and calculated the rlog using DESeq2. To calculate up- or down-regulation, we computed the rlog difference of the average of the triplicates against the baseline value (average over all conditions) for each gene.

### Western blot analysis

Western blot analysis for human iPSC–derived muscles was performed using antibodies to Dystrophin (NCL-DYS1, Leica), gamma-dystroglycan DAG1 (SC-165997, Santa Cruz Biotechnologies) and glyceraldehyde-3-phosphate dehydrogenase (MAB374, Millipore). Goat anti-mouse and goat anti-rabbit horseradish peroxidase–conjugated secondary antibodies were used for described experiments.

### Immunohistochemistry

Primary myogenic cultures were generated as described above. For secondary cultures, cells were replated on Matrigel (354277, Corning) coated glass bottom plates at a density of 52,000 cells/cm^2^. Cells were replated in SKGM medium (SKGM-2, Lonza CC-3245) supplemented with Rock inhibitor (#1254, R&D Systems). The next day, medium was replaced by fresh SKGM. Cells were then differentiated in KC/KCTi/KCTiP medium, cell cultures were fixed for 20 minutes in 4% paraformaldehyde (15710, Electron Microscopy Sciences) at room temperature. Cultures were rinsed three times in phosphate-buffered saline (PBS), followed by blocking buffer composed of PBS supplemented with 3% heat inactivated donkey serum (017-000-121, Jackson Immuno Research) and 0.1% Triton X-100 (T8787, Sigma-Aldrich). Primary antibodies were then diluted in blocking buffer and incubated overnight at 4°C. Cultures were then washed three times with PBS and incubated with secondary antibodies (Donkey Anti-Mouse/Rabbit IgG H+L Alexa Fluor® Cross-Adsorbed Secondary Antibody, 1:500) and Hoechst (5 μg/ml) in blocking buffer for 1 hour at room temperature. Cultures were then washed and stored in PBS until analyzed. Images were captured using a Zeiss LSM780 confocal microscope using 10x and 20x objectives. Images were analyzed using Fiji (Schindelin et al., 2012).

Primary antibodies used in this study:

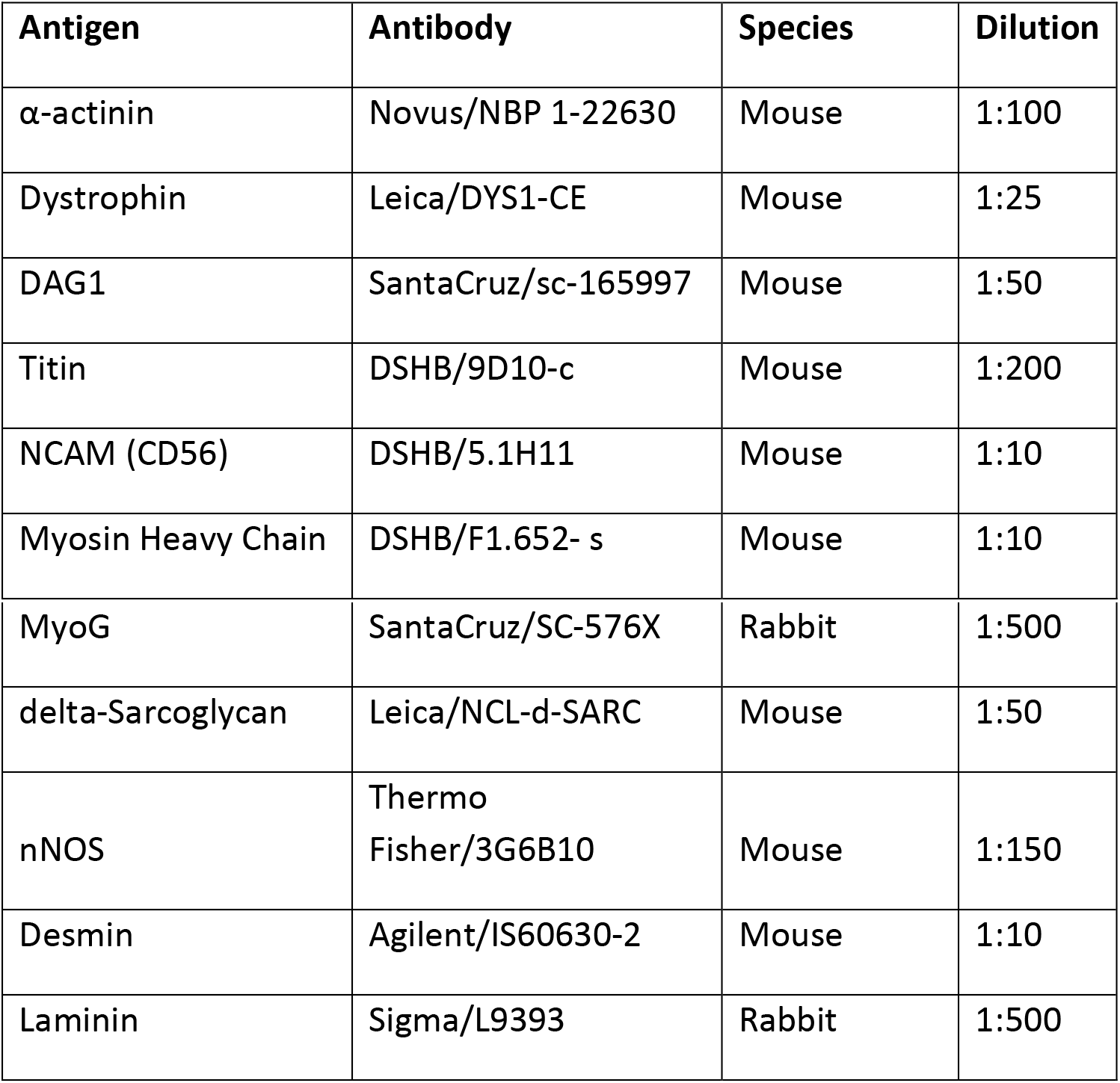

### Fiber branching quantification

Myogenic progenitors were replated in SKGM-2 medium supplemented with Rock inhibitor in 6-well plates. 24h after, cells were transfected with Tol2 CAGGS-nls mCherry IRES GFPcaax and CAGGS transposase (kindly donated by C. Marcelle, ARMI, Australia). Expression of the transposase enzyme along with flanking Tol2 sites ensure integration of the expression sequence and high fluorophore expression throughout expansion and differentiation. Plasmid-containing cells expressed ubiquitous NLS-mCherry in the nuclei and EGFP in the membrane. In this experimental set-up, any myofiber is immediately labelled upon fusion with a transfected cell. Cells were transfected with the plasmids using Lipofectamine 3000 (Thermofisher) at standard concentrations as suggested in the manufacturer protocols. This technique allowed us to permanently mark a subset of randomly-selected progenitors within the population, allowing for better visualization of individual fibers. Cells were allowed to recover 24 hours after lipofectamine, meaning differentiation was induced 48 h after replating.

One day after transfection (i.e. 48h after replating), cells were induced for differentiation using KC, KCTi, or KCTiP media. Cells were allowed to differentiate for 7 days, a sufficient time to allow for the formation of a large percentage of multinucleated fibers, as seen under an EVOS fluorescent microscope (Life Technologies).

Plates were then fixed with 4% PFA and immuno-stained for GFP (Abcam ab13970) and RFP (Abcam ab62341) and MF20 (DSHB) to enhance the fluorescent signal and confirm the identity of differentiated fibers, and the entire wells were imaged using the InCell 2000 arrayscan imaging platform (G.E. Life Sciences). Individual images (approximately 500 per well) were stitched using the Grid Stitching Plugin (Stephan Preibisch) on Fiji software. The number of branching points and total number of nuclei per fiber (all fibers visible from tip to tip were manually counted (approximately 500 fibers per well), fibers that could not be made out individually from others were not, each entire well was analyzed) were quantified. Averages and significance were statistically analyzed using a Kruskal-Wallis non-parametric ANOVA test with planned multiple comparisons (Prism 8 software, GraphPad).

### Contractility / Force measurements

#### Gelatin Muscular Thin Film Manufacture

Gelatin muscular thin film (MTF) chips were manufactured as previously reported (McCain et al., 2014), topaz or acrylic substrate surfaces were covered in a low adhesive tape (5560; Patco), and chips of 75×25 mm in dimensions were cut using a laser engraver. (Epilog Laser, Golden, CO). Tape was selectively peeled from each coverslip such that the base of each muscular thin film was revealed. Exposed regions were then oxygen plasma treated for one minute, (100 W RF, Plasma Prep II, SPI Supplies, West Chester, PA) to promote gelatin adhesion. Tape was then peeled from the remaining chip surface, exposing gelatin runoff channels. Onto each chip, 300 μL of 10% w/v gelatin was aliquoted. Quickly after aliquoting, a PDMS stamp containing aligned grooves of 20 μm x 20 μm (10 μm depth) was placed onto each chip. Next, on top of each stamp a glass slide to distribute mass and a 200 g weight were placed. Chips were then allowed to cure for a minimum of seven hours under moist conditions, at 4° C before being air dried and laser cut into thin film cantilevers, each 5×2 mm in dimension. Excess gelatin combs were then removed, and chips were adhered to the bottom of a 24-well petri dish using an additional 10% w/v gelatin glue. Chips were then stored in fresh phosphate buffered saline (PBS), and were UV sterilized prior to use. 10% w/v Gelatin solutions were prepared by dissolving 1.0 g of gelatin powder (Porcine Skin, Sigma-Aldrich) into 5 mL of deionized water, and 0.4 g of microbial transglutaminase (MTG) into another 5 mL of deionized water. Solutions were heated to 60° C and 37° C respectively until all powders dissolved (~30 minutes). Gelatin and MTG solutions were then combined and stirred using a vortex mixer. The resulting solution was then briefly degassed using a desiccator (~2 min), and the solution was then transferred to a 37° C water bath, where it was stored during MTF chip manufacture to prevent premature gelation.

#### Muscular Thin Film Contractile Experiments

Myogenic progenitors were generated from primary cultures of iPSC cells lines were differentiated for 20-30 days as described above and dissociated cells were replated on patterned gelatin MTFs in SKGM medium for 1-2 days to reach 80-90% of confluence. Cells were then differentiated over one week period in media containing Knock-out Replacement Serum (KSR), Chir 99021, and TGFβ inhibitor at 10 μM. Samples were treated with 10 μM Prednisolone during differentiation when indicated (KCTiP medium). After one week of differentiation muscular thin film experiments were performed in a Tyrode’s running solution (5 mM HEPES, 5.4 mM KCl, 1.8 mM CaCl2, 1 mM MgCl2, 135 mM NaCl, 5 mM glucose, and 0.33 mM NaH2PO4 in deionized water, pH 7.4; Sigma-Aldrich). After forming tissue constructs, the MTFs were peeled from the substrate, allowing them to contract and curl away from the underlying resting plane. Video micrographs were recorded on a Zeiss Discovery.V12 stereomicroscope using a Basler electric ACA2500-14UC USB 3.0 camera, at 30 frames per second, with a magnification of 7.2x and a resolution of 1920×1080 pixels. Thin films were recorded for an ~1.5 second period in the absence of stimulation, and then for 6.5 seconds at 1, 2, 10, 50 & 99hz at 20-30 V of stimulation. Field stimulation was controlled using an IonOptix myopacer with two parallel platinum electrode wires positioned ~30 cm apart. Each stimulation pulse was biphasic with a total duration of 10 ms per pulse, with 30 second breaks given between each stimulation time course.

Contractile stress was calculated using a custom python script. To measure contractile strength, the radius of curvature, *r*, was determined for each cantilever, using the following relation to numerically approximate *r*:

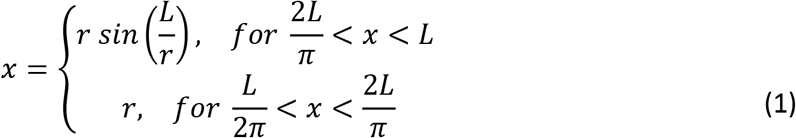

where, *L*, is the base cantilever length and *x,* is the x-projection of each cantilever. X-projections were extracted from cantilever movies using a custom script in NIH’s ImageJ. Briefly, regions of interest were determined along the midsection for each cantilever, running from the base of the MTF to the inscribed edge. Each cantilever image was then thresholded to a binary and x-projections were extracted. The radius of curvatures was then converted to contractile stresses, σ_c_, using a modified version of Stoney’s equation for deforming thin sheets, such that:

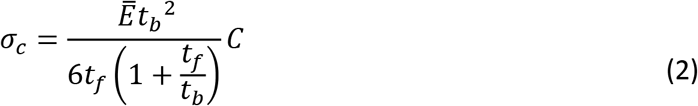

Where 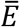 is the film’s Youngs modulus in the direction of contraction, *t_b_* is the gelatin film’s thickness, *tf* is the tissue thickness, and c is the film curvature as given by 1/*r*. Here an average film thickness of 134 μm and a tissue thickness of 15 μm were used (as determined by confocal microscopy, supplementary Figure 6). A young’s modulus of 55.6 kPA was used for gelatin films as has been previously reported for this protocol (McCain et al., 2014).

Contractile stress here is reported as the difference between maximal contraction, and the baseline pre-strain, as determined on a per cantilever basis. As the stimulation protocols were manually initiated, contraction time courses were corrected by using the maximal value of the first derivative of contraction vs time to match stimulation times. Contractile time course measurements were additional internally normalized by subtracting the minimum recorded prestrain from each cantilever measurement.

#### Specific Tensile Strength Estimates

To normalize thin film contractile stress against *in vivo* tensile models, we estimated the specific tensile strength, *T_s_*, of the engineered tissue constructs using dimensional analysis such that:

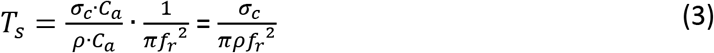

Where *f_r_* is the fiber bundle radius which we used to approximate the physiological cross-section, *σ_c_*was the maximal contractile stress, *C_a_* is the cantilever area and ρ is the fiber density. To estimate fiber density and cross-sectional areas, we used orthogonal confocal fluorescent cross sections (Supplementary Figure 6) of aligned tissue. This yielded an estimate of ~620±280 muscle fibers per cantilever. Uncertainties in tensile strengths are reported as the standard error of the mean.

### Calcium signaling analysis

#### Cell Culture and Sample Preparation

Human iPSC-derived myoblasts were differentiated into myocyte fibers according to (Chal et al., 2016). 3-4 weeks old primary myogenic cultures generated from wild-type iPSCs were dissociated as described and myogenic progenitors (myoblasts) were replated at low density (35-40k/cm^2^) onto Matrigel (Corning, Cat#354277)-coated dishes in skeletal muscle growth media (SKGM-2, Lonza CC-3245) with 10 μM ROCK inhibitor. After 24 hours, medium was changed to SKGM-2 media without ROCK inhibitor and incubated with low-titer lentivirus encoding CheRiff-CFP. Myoblast cultures were allowed to proliferate for up to 72 hours, at which point cultures reached ~90% confluence. Cultures were then induced for myogenic differentiation with DMEM/F12 supplemented with 2% knock-out serum replacement (Invitrogen, Cat. # 10828028), 10 μM of the TGFβ inhibitor SB431542 (Tocris, Cat. # 1614), 1 μM Chiron (Tocris, Cat. # 4423), 0.2% Pen/Strep (Life Technologies, Cat. # 15140122) and 1x ITS (Life Technologies, Cat. # 41400045). Following induction, medium was changed on days 1 and 2 and then was refreshed every other day for up to 10 days post-differentiation to generate mature and fused myocyte fibers.

CheRiff-expressing myocyte cultures were stained for imaging after 7 days of differentiation (10 days total culture time). Samples were stained either with CaSiR-1 AM (Goryo Chemical) or BioTracker 609 AM Red Ca^2+^ Dye (EMD Millipore). Briefly, on the day of staining, 50 μg dye aliquots were thawed and then reconstituted in DMSO to 1 mM stock concentration. A loading solution was prepared by diluting calcium dye stocks 1:500 in phosphate buffered saline to 2 μM concentration in the presence of 0.02% final concentration of Pluronic F127 (Sigma). Myofiber samples were stained in loading solution for 30 minutes in a standard tissue-culture incubator, washed twice in phosphate buffered saline to remove residual dye, and then transferred to Tyrode’s solution containing (in mM) 125 NaCl, 2 KCl, 2 CaCl2, 1 MgCl2, 10 HEPES, 30 glucose. The pH was adjusted to 7.3 with NaOH and the osmolality was adjusted to 305–310 mOsm with sucrose. All calcium measurements were performed in 2 mL of Tyrode’s solution buffer.

#### Optogenetic Stimulation and Calcium Imaging

Spatially resolved optical electrophysiology measurements were performed using a home-built upright ultra-widefield microscope (Werley et al., 2017) with a large field of view (4.6×4.6 mm^2^, with 2.25×2.25 μm^2^ pixel size) and high numerical aperture objective lens (Olympus MVPLAPO 2XC, NA 0.5). Fluorescence of CaSiR-1 was excited with a 639 nm laser (OptoEngine MLL-FN-639) at 100 mW/cm^2^, illuminating the sample from below at an oblique angle to minimize background autofluorescence. CaSiR-1 fluorescence was separated from scattered laser excitation via a dichroic beam splitter (Semrock Di01-R405/488/561/635-t3-60×85) and an emission filter (Semrock FF01-708/75-60-D). BioTracker 609 fluorescence was excited using 561 nm laser beam (MPB Communications, F-04306-02) with the same dichroic beam splitter with a separate emission filter (Chroma ET600/50m). Images were collected at a 50 Hz frame rate on a Hamamatsu Orca Flash 4.2 scientific CMOS camera. Optogenetic stimulation was performed by exciting CheRiff with a blue LED (Thorlabs M470L3) with a fixed intensity between 10 and 30 mW/cm^2^ and patterned using a digital micromirror device module (Vialux V-7001). For each experiment, blue illumination intensity was first adjusted relative to CheRiff expression in a test sample dish in order to elicit a range of calcium responses across frequencies, and then fixed for all samples for the remainder of the experiment.

#### Image Processing and Data Analysis

Optical recordings of calcium-sensitive CaSiR-1 AM and BioTracker 609 AM fluorescence were processed using custom MATLAB software. Briefly, to minimize uncorrelated shot-noise, movies were subjected to 4×4 binning. Changes in fluorescence were then calculated relative to an initial baseline image in order to calculate the calcium-sensitive component of the signal (ΔF/F). ΔF/F movies were then averaged across the optical stimulation region of interest to generate an overall sample response, which was then median filtered in the time domain (9 frame kernel). Individual dish responses were then normalized relative to the response to a saturating channelrhodopsin stimulus in order to account for differences in labeling density. For ionophore-normalized controls, responses were instead normalized relative to the ΔF/F response to bath application of ionomycin (10 μM). For BioTracker 609 AM data, an additional crosstalk subtraction step was performed for 20 Hz stimulation periods due to unfiltered bleedthrough of fluorescence excited by the 470 nm stimulus.

Statistical significance between cell lines was assessed via paired two-sample t-tests using standard MATLAB functions (ttest2). For a given sample dish, calcium responses were averaged within a particular stimulus train in order to calculate an overall response metric per dish per frequency, and then populations of responses were compared at each frequency between relevant experimental conditions (e.g. NCRM1 vs DMDI vs DMDII; TX1-COR vs TX1-UNC). Figures indicate confidence intervals of p<0.05 (*), p<0.01(**), and p<0.001(***). For isogenic lines, N=6 samples were compared for each condition (NCRM1, DMDI, DMDII) for 18 total measurements. For patient-derived cells, N=6 samples were analyzed for both TX1-COR and TX1-UNC for the first replicate, and N=4 samples were analyzed for each condition in the second replicate (comprising 20 total patient-derived dishes measured).

## Supplementary Tables

**Supplementary Table 1:** List of genes upregulated or downregulated (DESEQ) in DMDI and DMDII vs WT NCRM1 iPSC cells isolated after a week of secondary differentiation in KC medium

**Supplementary Table 2:** EnrichR analysis of the genes upregulated and downregulated in DMDI and DMDII vs WT NCRM1 iPSC cells isolated after a week of secondary differentiation in KC medium

**Supplementary Table 3:** Number of isogenic MTF cantilevers assayed as a function of stimulation frequency. Number of cantilevers gives the amount of individual force measurements recorded for each condition (3-4 cantilevers per well), while number of chip gives the number of distinct replicates.

**Supplementary Table 4:** Number of patient derived MTF cantilevers assayed as a function of stimulation frequency. Number of cantilevers gives the amount of individual force measurements recorded for each condition (3-4 cantilever’s per well), while number of chip gives the number of distinct replicates.

**Supplementary Table 5:** Guide RNAs used for the construction of the DMDI and DMDII lines and primers used for clone screening

## Supplementary Movies

**Supplementary Movie 1: Contractile phenotype of isogenic hIPSC derived skeletal muscle.** Representative brightfield micrograph of muscular thin films seeded with DMD (DMD II - upper, left) (DMD I-lower left) and wildtype (wt) cell lines (upper, right), in the absence (left-KCTi) and presence (right-KCTiP) of prednisolone, with corresponding force measurements (lower right) (scale bars – 1 mm) (99 Hz stimulation induced at ~2 s). Uncertainty in force measurements given as the standard error in measurement for the whole chip (full chip not shown).

**Supplementary Movie 2: Contractile phenotype of patient derived skeletal muscle.** Representative brightfield micrograph of muscular thin films seeded with patient derived uncorrected DMD (UNC - lower, left & upper, right) cells and CAS9 corrected cell lines (Cor lowerr, right), in the absence (right-KCTi) and presence (left) of prednisolone, with corresponding force measurements (upper left) (scale bars – 1 mm) (99 Hz stimulation induced at ~2 s). Uncertainty in force measurements given as the standard error in measurement.

**Supplementary Movie 3: Calcium responses in isogenic cell lines under optogenetic drive.** NCRM1, DMDI, and DMDII cells were inoculated with lentivirus encoding the channelrhodopsin CheRiff, and then differentiated in parallel. After 7 days in differentiation media, cells were stained with the calcium sensitive dye CaSiR-1-AM (Goryo Chemical), and then transferred to a home-built microscope to image calcium responses to optogenetic stimulation. The movie shows a montage of calcium responses from each of 3 representative dishes (1 each from the 3 experimental conditions). Samples were stimulated with a patterned pulses (20 ms) of 470 nm light in the central region, at a 0.5 Hz frequency (denoted by cyan overlays). Calcium responses are displayed in units of deltaF/F (see Methods) and pseudocolored according to the MATLAB colormap ‘hot’ (0 to 25% dF/F dynamic range). Movies were acquired at 20 Hz and are played back at 10 fps (i.e. 0.5x real-time speed).

**Supplementary Movie 4: Calcium responses in patient-derived cells under optogenetic drive.** Uncorrected (TX1-UNC) and corrected (TX1-COR) varients of a patient-derived cell line were innoculated with lentivirus encoding the channelrhodopsin CheRiff, and then differentiated in parallel. After 7 days in differentiation media, cells were stained with the calcium sensitive dye CaSiR-1-AM (Goryo Chemical), and then transferred to a home-built microscope to image calcium responses to optogenetic stimulation. The movie shows a montage of calcium responses from both corrected and uncorrected cells in parallel. Samples were stimulated with a patterned pulses (20 ms) of 470 nm light in the central region, at a 0.5 Hz frequency (denoted by cyan overlays). Calcium responses are displayed in units of deltaF/F (see Methods) and pseudocolored according to the MATLAB colormap ‘hot’ (0 to 25% dF/F dynamic range). Movies were acquired at 50 Hz and are played back at 25 fps (i.e. 0.5x real-time speed).

**Supplementary Movie 5: Calcium responses in patient-derived cells under optogenetic drive, replicated.** An independent set patient-derived cells was performed to confirm the results in Movie 2. As before, differentiations of uncorrected (TX1-UNC) and corrected (TX1-COR) variants of a patient-derived cell line were innoculated with lentivirus encoding the channelrhodopsin CheRiff, and then differentiated in parallel. After 7 days in differentiation media, cells were stained with the calcium sensitive dye BioTracker 609 AM (Sigma), and then transferred to a home-built microscope to image calcium responses to optogenetic stimulation. The movie shows a montage calcium responses from both corrected and uncorrected cells in parallel. Samples were stimulated with 50 ms pulses of widefield 470 nm light at a 0.5 Hz frequency (denoted by cyan overlays). Calcium responses are displayed in units of deltaF/F (see Methods) and pseudocolored according to the MATLAB colormap ‘hot’ (0 to 25% dF/F dynamic range). Movies were acquired at 50 Hz, and are played back at 25 fps (i.e. 0.5x real-time speed).

## Supplementary Figures

**Supplementary Figure 1:**
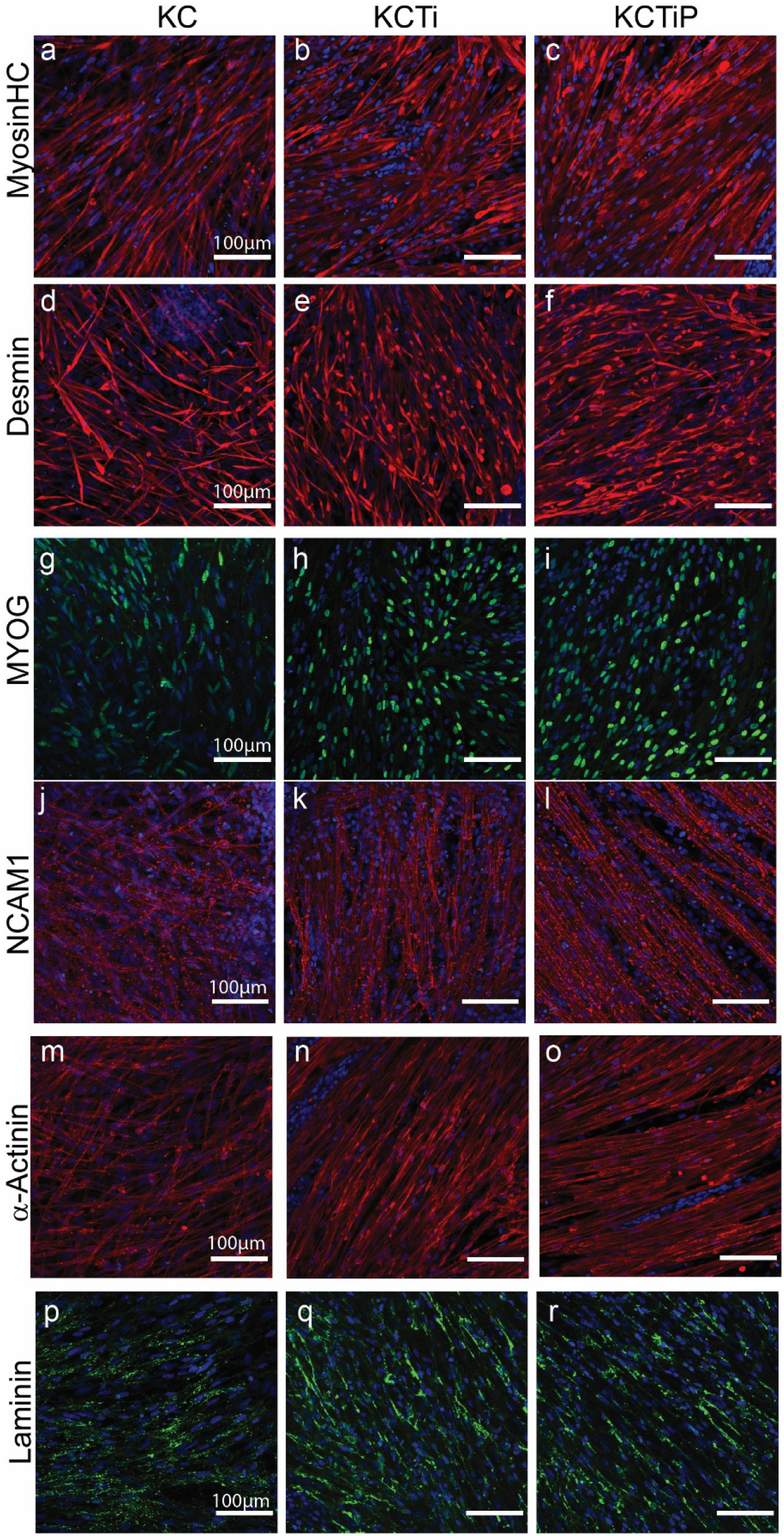
Comparison of KC, KCTi and KCTiP differentiation conditions. Immunocytochemistry analysis of the wild-type iPSC line differentiated in KC, KCTi or KCTiP medium for one week. (a-c) MyosinHC, Myosin heavy chain (d-f) Desmin (g-i) Myogenin (MYOG) (j-l) Neural Cell Adhesion Molecule 1 (NCAM1) (m-o), α-actinin (p-r) Laminin. Note increased fiber thickness and alignment in KCTiP medium.

**Supplementary Figure 2:**
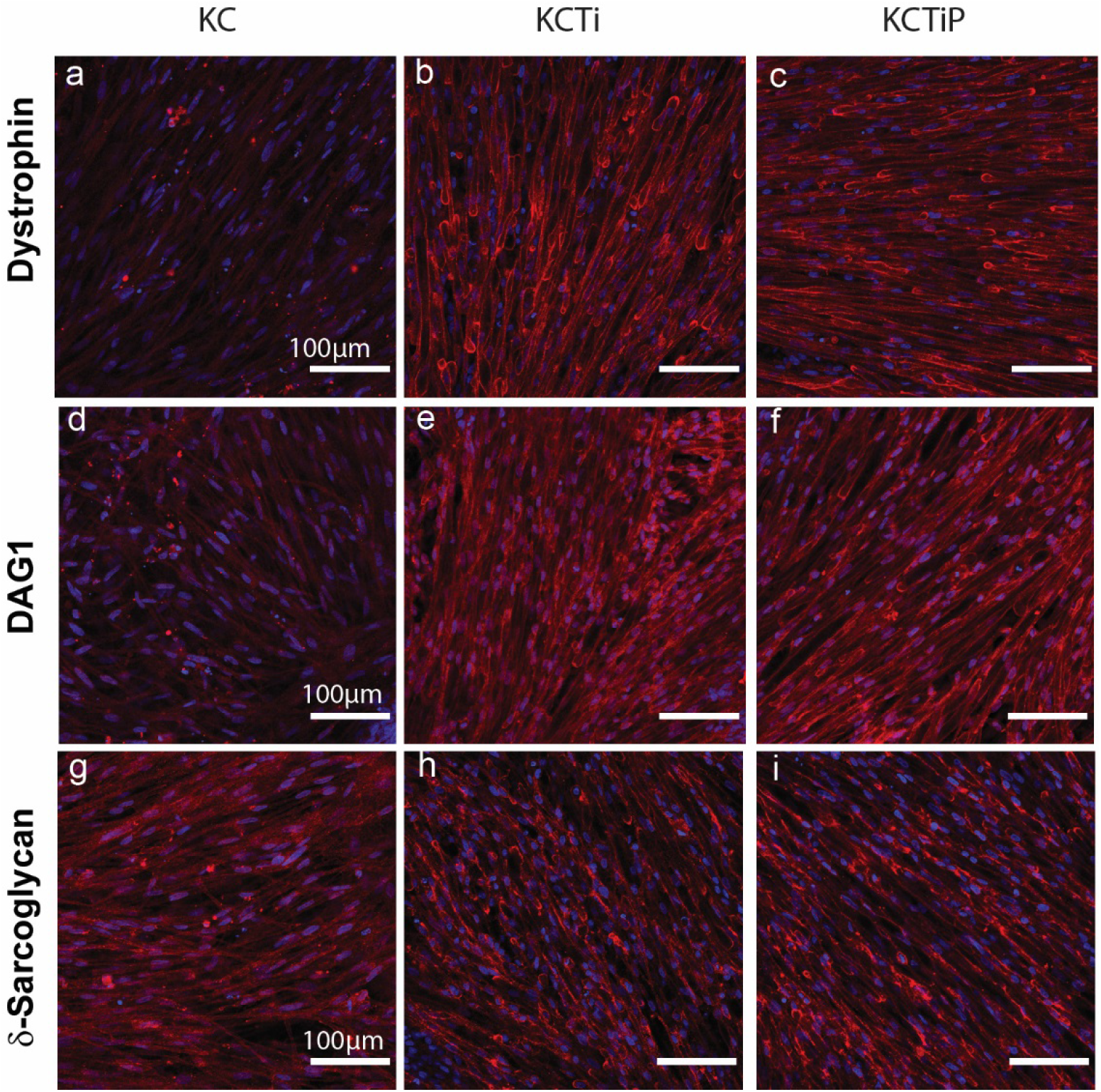
Comparison of DGC proteins expression in KC, KCTi and KCTiP differentiation conditions. Immunocytochemistry of DGC genes Dystrophin (a-c), DAG1 (d-f) and delta-Sarcoglycan (g-h) in the NCRM1 iPSC line after one week of secondary differentiation in KC, KCTi, and KCTiP media. Note increased expression of the three proteins in KCTi and KCTiP medium and changes in delta-Sarcoglycan localization and clustering.

**Supplementary Figure 3:**
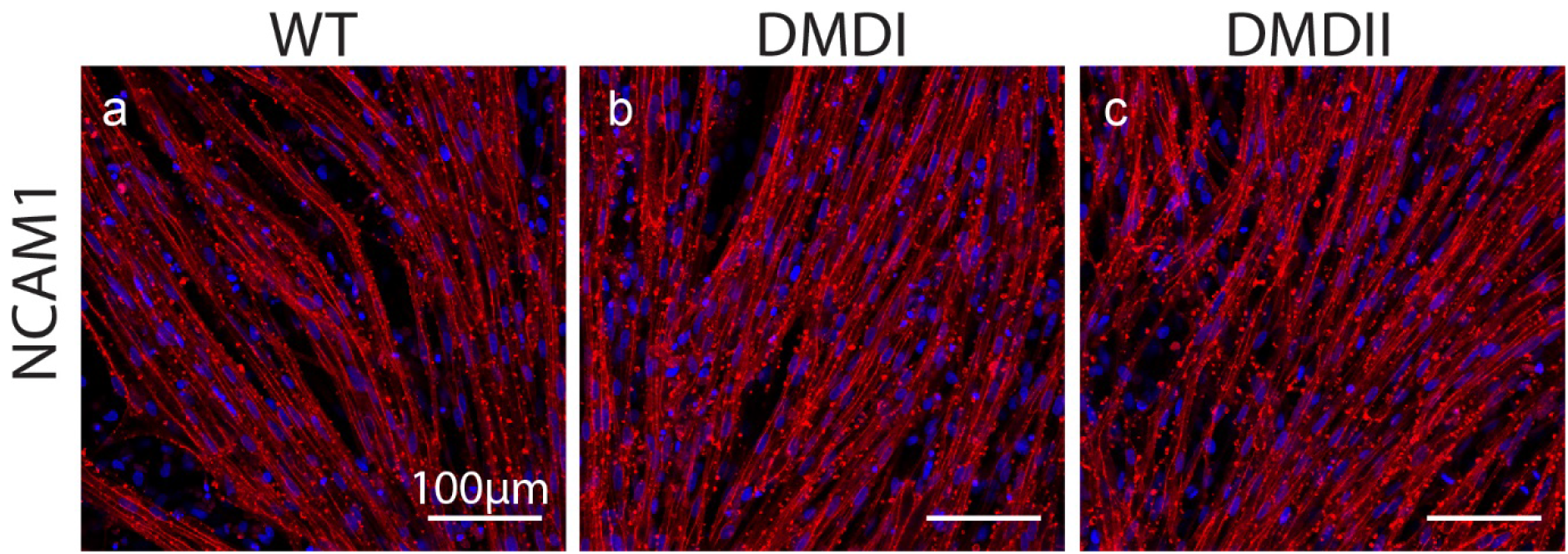
Immunocytochemistry using anti-Neural Cell Adhesion Molecule 1 (NCAM1) after one week of secondary differentiation in KCTiP medium. (a) WT = wild-type NCRM1 line, (b) DMDI, (c) DMDII – DMD deficient isogenic lines.

**Supplementary Figure 4:**
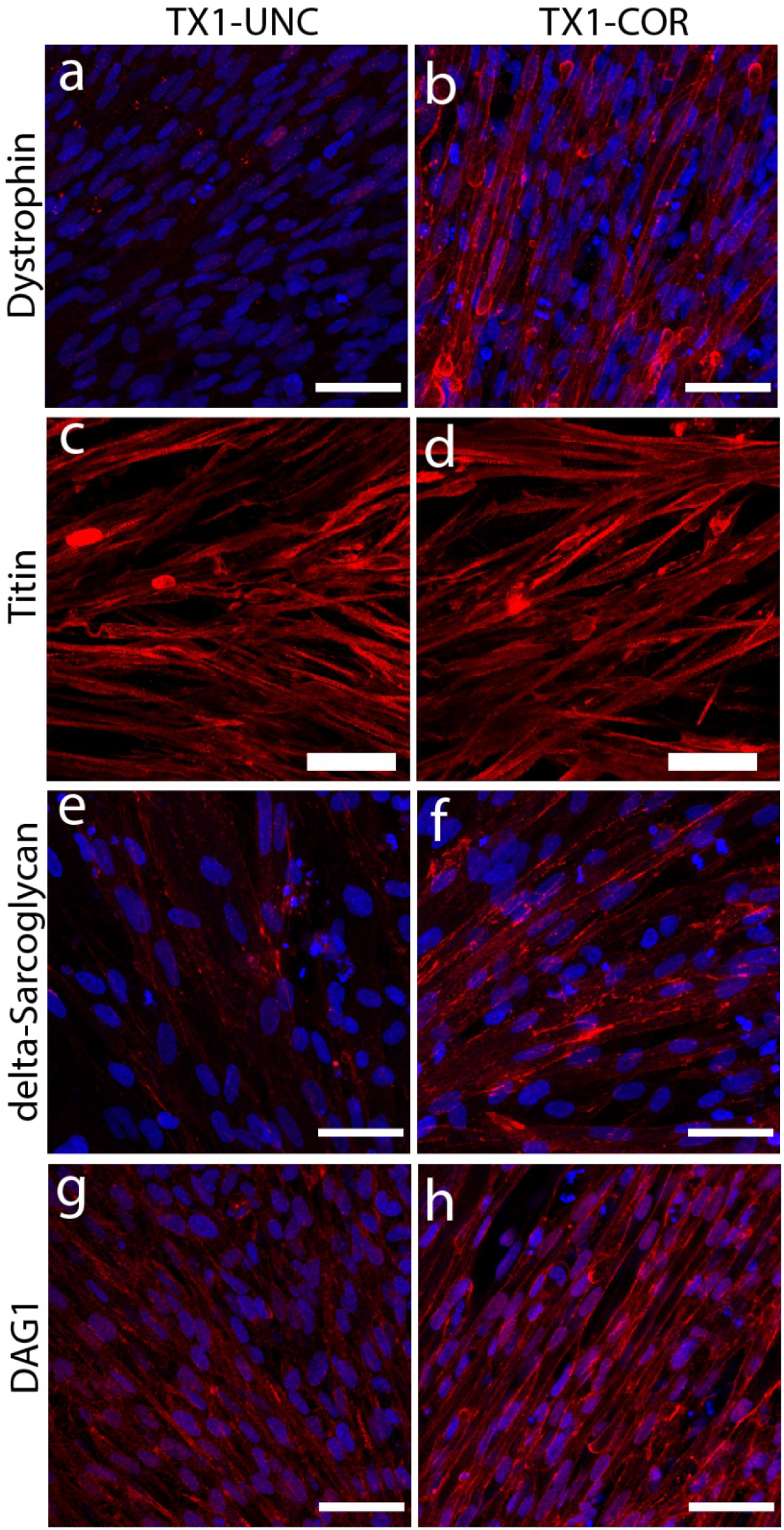
Immunocytochemistry analysis using antibodies for Dystrophin (a, b), Titin (c, d), delta-sarcoglycan (e, f), and DAG1 (g,h) in TX1-UNC and TX1-COR cell lines after one week of secondary differentiation in KCTiP medium. Note the absence of Dystrophin, decrease in delta-sarcoglycan, and DAG1 antibody signal in TX1-UNC cells.

**Supplementary Figure 5:**
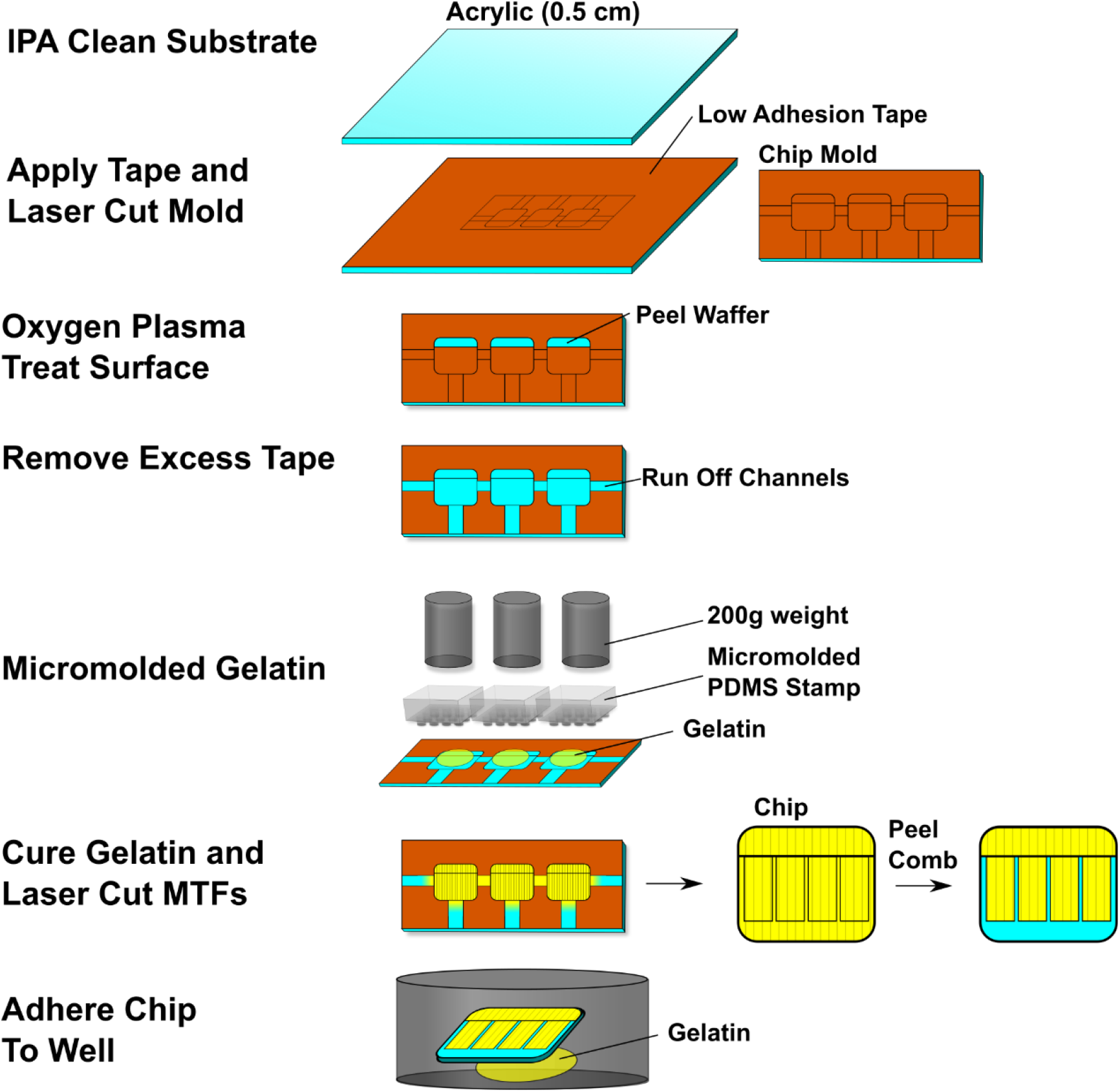
Cantilever Chip Preparation Flow Chart. Muscle Thin Films (MTF) chips are first prepared with a low adhesion tape, and then laser cut into coverslip sized regions. The low adhesive tape is then removed from selected regions, to allow surface treatment, causing the gelatin to adhere to the exposed base. Run off channels are then removed, and gelatin is added, which is then compressed using a micromolded PDMS stamp with an additional 200g weight. Individual chips are then laser cut out, and adhered to the bottom of a well plate to facilitate cell culture. Before use, all samples are UV sterilized.

**Supplementary Figure 6.**
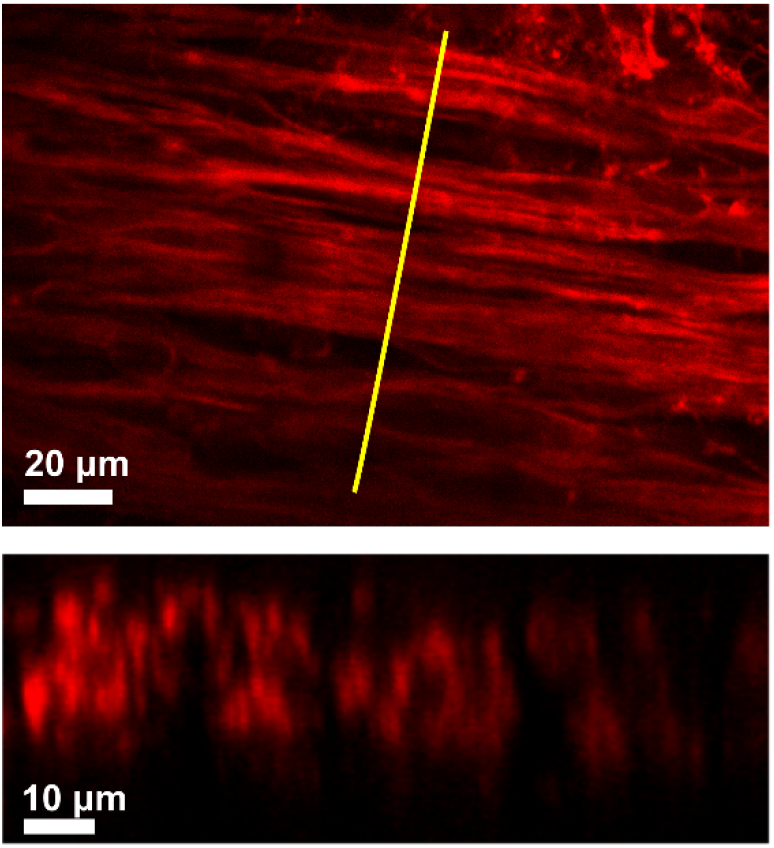
Aligned isogenic cell cross section. Representative confocal fluorescent micrograph (upper) of aligned wild-type isogenic cells after culture on gelatin substrates, with corresponding orthogonal cross section (lower) taken from the highlighted region showing myofibril bundles cross sections (red-actin).

**Supplementary Figure 7:**
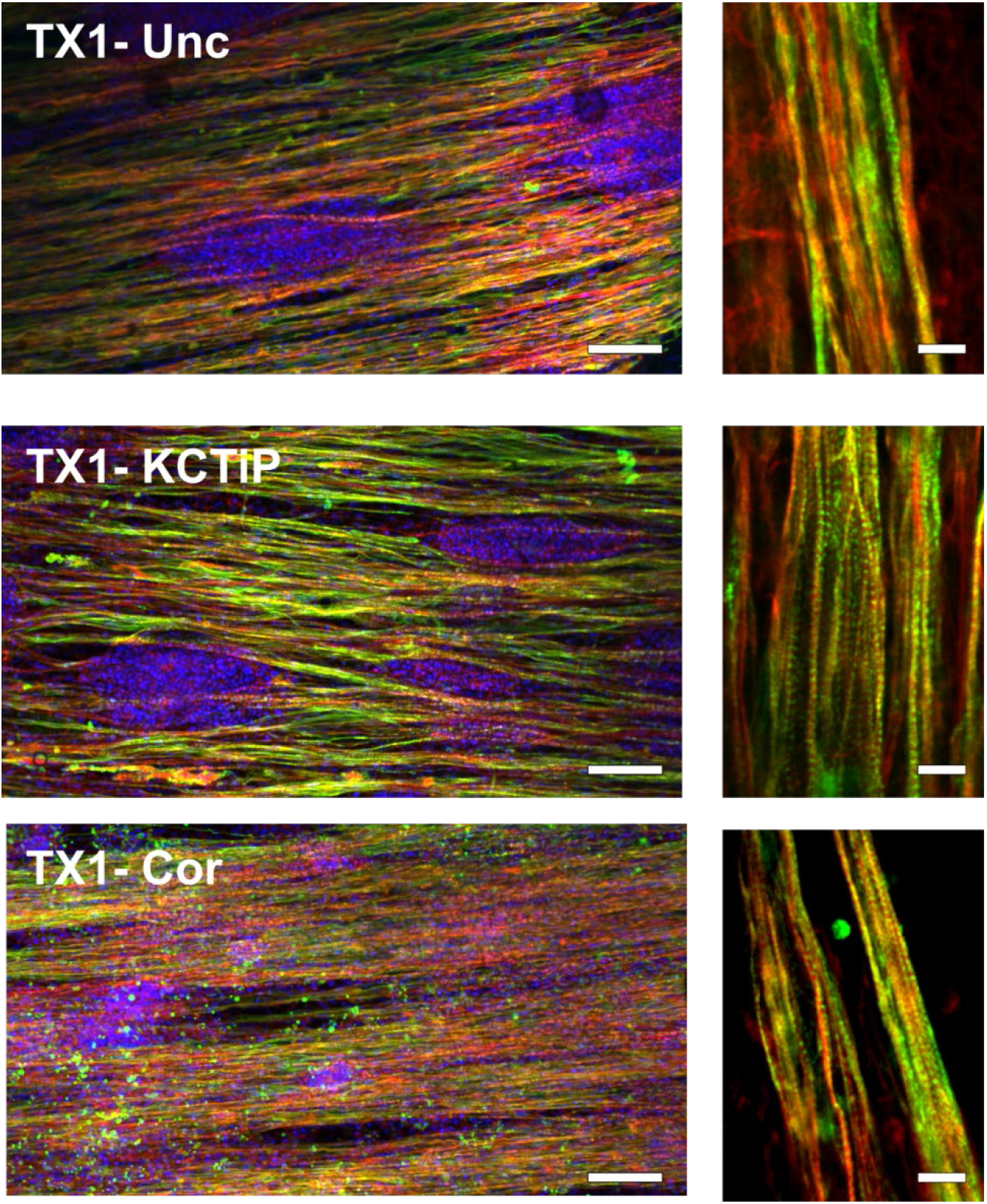
Representative immunofluorescence micrograph of patient derived skeletal muscles grown on micro-molded gelatin substrates. Showing confluent aligned tissues (Uncorrected, TX1-Unc – Upper, Prednisolone treated, KCTiP – Middle, Cas9 corrected to healthy phenotype, TX1-Cor – Lower) (DAPI: Blue, α-actinin: green, actin: Red), with high magnification inset showing sarcomere expression (Scale bars: 100 μm, left; 10 μm, right)

## References

Al Tanoury, Z., Rao, J., Tassy, O., Gobert, B., Gapon, S., Garnier, J.M., Wagner, E., Hick, A., Hall, A., Gussoni, E., et al. (2020). Differentiation of the human PAX7-positive myogenic precursors/satellite cell lineage in vitro. Development.

Braun, S., Tranchant, C., Vilquin, J.T., Labouret, P., Warter, J.M., and Poindron, P. (1989). Stimulating effects of prednisolone on acetylcholine receptor expression and myogenesis in primary culture of newborn rat muscle cells. J Neurol Sci 92, 119.–131.

Brenman, J.E., Chao, D.S., Xia, H., Aldape, K., and Bredt, D.S. (1995). Nitric oxide synthase complexed with dystrophin and absent from skeletal muscle sarcolemma in Duchenne muscular dystrophy. Cell 82, 743–752.

Burr, A.R., and Molkentin, J.D. (2015). Genetic evidence in the mouse solidifies the calcium hypothesis of myofiber death in muscular dystrophy. Cell death and differentiation 22, 1402–1412.

Busada, J.T., and Cidlowski, J.A. (2017). Mechanisms of Glucocorticoid Action During Development. Curr Top Dev Biol 125, 147–170.

Chal, J., Al Tanoury, Z., Hestin, M., Gobert, B., Aivio, S., Hick, A., Cherrier, T., Nesmith, A.P., Parker, K.K., and Pourquie, O. (2016). Generation of human muscle fibers and satellite-like cells from human pluripotent stem cells in vitro. Nature protocols 11, 1833–1850.

Chal, J., Oginuma, M., Al Tanoury, Z., Gobert, B., Sumara, O., Hick, A., Bousson, F., Zidouni, Y., Mursch, C., Moncuquet, P., et al. (2015). Differentiation of pluripotent stem cells to muscle fiber to model Duchenne muscular dystrophy. Nat Biotechnol 33, 962–969.

Chal, J., and Pourquie, O. (2017). Making muscle: skeletal myogenesis in vivo and in vitro. Development 144, 2104–2122.

Chan, S., and Head, S.I. (2011). The role of branched fibres in the pathogenesis of Duchenne muscular dystrophy. Experimental physiology 96, 564–571.

Choi, I.Y., Lim, H., Estrellas, K., Mula, J., Cohen, T.V., Zhang, Y., Donnelly, C.J., Richard, J.P., Kim, Y.J., Kim, H., et al. (2016). Concordant but Varied Phenotypes among Duchenne Muscular Dystrophy Patient-Specific Myoblasts Derived using a Human iPSC-Based Model. Cell reports 15, 2301–2312.

Costa, P.B., Ryan, E.D., Herda, T.J., Defreitas, J.M., Beck, T.W., and Cramer, J.T. (2009). Effects of static stretching on the hamstrings-to-quadriceps ratio and electromyographic amplitude in men. J Sports Med Phys Fitness 49, 401–409.

Diaz-Cuadros, M., Wagner, D.E., Budjan, C., Hubaud, A., Tarazona, O.A., Donelly, S., Michaut, A., Al Tanoury, Z., Yoshioka-Kobayashi, K., Niino, Y., et al. (2020). In vitro characterization of the human segmentation clock. Nature 580, 113–118.

Emery, A.E. (1977). Muscle histology and creatine kinase levels in the foetus in Duchenne muscular dystrophy. Nature 266, 472–473.

Emery, A.E., Burt, D., Dubowitz, V., Rocker, I., Donnai, D., Harris, R., and Donnai, P. (1979). Antenatal diagnosis of Duchenne muscular dystrophy. Lancet 1, 847–849.

Esposito, G., Tremolaterra, M.R., Marsocci, E., Tandurella, I.C., Fioretti, T., Savarese, M., and Carsana, A. (2017). Precise mapping of 17 deletion breakpoints within the central hotspot deletion region (introns 50 and 51) of the DMD gene. J Hum Genet 62, 1057–1063.

Farini, A., Sitzia, C., Cassinelli, L., Colleoni, F., Parolini, D., Giovanella, U., Maciotta, S., Colombo, A., Meregalli, M., and Torrente, Y. (2016). Inositol 1,4,5-trisphosphate (IP3)-dependent Ca2+ signaling mediates delayed myogenesis in Duchenne muscular dystrophy fetal muscle. Development 143, 658–669.

Fink, R.H., Stephenson, D.G., and Williams, D.A. (1990). Physiological properties of skinned fibres from normal and dystrophic (Duchenne) human muscle activated by Ca2+ and Sr2+. J Physiol 420, 337–353.

Fong, P.Y., Turner, P.R., Denetclaw, W.F., and Steinhardt, R.A. (1990). Increased activity of calcium leak channels in myotubes of Duchenne human and mdx mouse origin. Science 250, 673–676.

Franco, A., Jr., and Lansman, J.B. (1990). Calcium entry through stretch-inactivated ion channels in mdx myotubes. Nature 344, 670–673.

Hicks, M.R., Hiserodt, J., Paras, K., Fujiwara, W., Eskin, A., Jan, M., Xi, H., Young, C.S., Evseenko, D., Nelson, S.F., et al. (2018). ERBB3 and NGFR mark a distinct skeletal muscle progenitor cell in human development and hPSCs. Nat Cell Biol 20, 46–57.

Hochbaum, D.R., Zhao, Y., Farhi, S.L., Klapoetke, N., Werley, C.A., Kapoor, V., Zou, P., Kralj, J.M., Maclaurin, D., Smedemark-Margulies, N., et al. (2014). All-optical electrophysiology in mammalian neurons using engineered microbial rhodopsins. Nature methods 11, 825–833.

Janghra, N., Morgan, J.E., Sewry, C.A., Wilson, F.X., Davies, K.E., Muntoni, F., and Tinsley, J. (2016). Correlation of Utrophin Levels with the Dystrophin Protein Complex and Muscle Fibre Regeneration in Duchenne and Becker Muscular Dystrophy Muscle Biopsies. PLoS ONE 11, e0150818.

Kanda, F., Okuda, S., Matsushita, T., Takatani, K., Kimura, K.I., and Chihara, K. (2001). Steroid myopathy: pathogenesis and effects of growth hormone and insulin-like growth factor-I administration. Horm Res 56 Suppl 1, 24–28.

Kissel, J.T., Burrow, K.L., Rammohan, K.W., and Mendell, J.R. (1991). Mononuclear cell analysis of muscle biopsies in prednisone-treated and untreated Duchenne muscular dystrophy. CIDD Study Group. Neurology 41, 667–672.

Long, C., Li, H., Tiburcy, M., Rodriguez-Caycedo, C., Kyrychenko, V., Zhou, H., Zhang, Y., Min, Y.L., Shelton, J.M., Mammen, P.P.A., et al. (2018). Correction of diverse muscular dystrophy mutations in human engineered heart muscle by single-site genome editing. Sci Adv 4, eaap9004.

Lowe, D.A., Williams, B.O., Thomas, D.D., and Grange, R.W. (2006). Molecular and cellular contractile dysfunction of dystrophic muscle from young mice. Muscle Nerve 34, 92–100.

Maganaris, C.N., Baltzopoulos, V., Ball, D., and Sargeant, A.J. (2001). In vivo specific tension of human skeletal muscle. Journal of applied physiology 90, 865–872.

Matthews, E., Brassington, R., Kuntzer, T., Jichi, F., and Manzur, A.Y. (2016). Corticosteroids for the treatment of Duchenne muscular dystrophy. Cochrane Database Syst Rev, CD003725.

McGreevy, J.W., Hakim, C.H., McIntosh, M.A., and Duan, D. (2015). Animal models of Duchenne muscular dystrophy: from basic mechanisms to gene therapy. Dis Model Mech 8, 195–213.

Merrick, D., Stadler, L.K., Larner, D., and Smith, J. (2009). Muscular dystrophy begins early in embryonic development deriving from stem cell loss and disrupted skeletal muscle formation. Dis Model Mech 2, 374–388.

Messina, G., Biressi, S., Monteverde, S., Magli, A., Cassano, M., Perani, L., Roncaglia, E., Tagliafico, E., Starnes, L., Campbell, C.E., et al. (2010). Nfix regulates fetal-specific transcription in developing skeletal muscle. Cell 140, 554–566.

Moretti, A., Fonteyne, L., Giesert, F., Hoppmann, P., Meier, A.B., Bozoglu, T., Baehr, A., Schneider, C.M., Sinnecker, D., Klett, K., et al. (2020). Somatic gene editing ameliorates skeletal and cardiac muscle failure in pig and human models of Duchenne muscular dystrophy. Nat Med 26, 207–214.

Nesmith, A.P., Wagner, M.A., Pasqualini, F.S., O’Connor, B.B., Pincus, M.J., August, P.R., and Parker, K.K. (2016). A human in vitro model of Duchenne muscular dystrophy muscle formation and contractility. J Cell Biol 215, 47–56.

Osafune, K., Caron, L., Borowiak, M., Martinez, R.J., Fitz-Gerald, C.S., Sato, Y., Cowan, C.A., Chien, K.R., and Melton, D.A. (2008). Marked differences in differentiation propensity among human embryonic stem cell lines. Nat Biotechnol 26, 313–315.

Partridge, T.A. (2013). The mdx mouse model as a surrogate for Duchenne muscular dystrophy. FEBS J 280, 4177–4186.

Pearson, C.M. (1962). Histopathological features of muscle in the preclinical stages of muscular dystrophy. Brain 85, 109–120.

Petrof, B.J., Shrager, J.B., Stedman, H.H., Kelly, A.M., and Sweeney, H.L. (1993). Dystrophin protects the sarcolemma from stresses developed during muscle contraction. Proc Natl Acad Sci U S A 90, 3710–3714.

Piga, D., Salani, S., Magri, F., Brusa, R., Mauri, E., Comi, G.P., Bresolin, N., and Corti, S. (2019). Human induced pluripotent stem cell models for the study and treatment of Duchenne and Becker muscular dystrophies. Ther Adv Neurol Disord 12, 1756286419833478.

Quattrocelli, M., Zelikovich, A.S., Jiang, Z., Peek, C.B., Demonbreun, A.R., Kuntz, N.L., Barish, G.D., Haldar, S.M., Bass, J., and McNally, E.M. (2019). Pulsed glucocorticoids enhance dystrophic muscle performance through epigenetic-metabolic reprogramming. JCI Insight 4.

Rahimov, F., and Kunkel, L.M. (2013). The cell biology of disease: cellular and molecular mechanisms underlying muscular dystrophy. J Cell Biol 201, 499–510.

Schiaffino, S., Rossi, A.C., Smerdu, V., Leinwand, L.A., and Reggiani, C. (2015). Developmental myosins: expression patterns and functional significance. Skeletal muscle 5, 22.

Schmalbruch, H. (1984). Regenerated muscle fibers in Duchenne muscular dystrophy: a serial section study. Neurology 34, 60–65.

Shoji, E., Sakurai, H., Nishino, T., Nakahata, T., Heike, T., Awaya, T., Fujii, N., Manabe, Y., Matsuo, M., and Sehara-Fujisawa, A. (2015). Early pathogenesis of Duchenne muscular dystrophy modelled in patient-derived human induced pluripotent stem cells. Sci Rep 5, 12831.

Sieiro, D., Melendez, J., Morin, V., Salgado, D., and Marcelle, C. (2019). Auto-inhibition of myoblast fusion by cyclic receptor signalling. Bioarxiv.

Sklar, R.M., and Brown, R.H., Jr. (1991). Methylprednisolone increases dystrophin levels by inhibiting myotube death during myogenesis of normal human muscle in vitro. J Neurol Sci 101, 73–81.

Voit, A., Patel, V., Pachon, R., Shah, V., Bakhutma, M., Kohlbrenner, E., McArdle, J.J., Dell’Italia, L.J., Mendell, J.R., Xie, L.H., et al. (2017). Reducing sarcolipin expression mitigates Duchenne muscular dystrophy and associated cardiomyopathy in mice. Nature communications 8, 1068.

Wang, G., McCain, M.L., Yang, L., He, A., Pasqualini, F.S., Agarwal, A., Yuan, H., Jiang, D., Zhang, D., Zangi, L., et al. (2014). Modeling the mitochondrial cardiomyopathy of Barth syndrome with induced pluripotent stem cell and heart-on-chip technologies. Nat Med 20, 616–623.

Wehling-Henricks, M., Lee, J.J., and Tidball, J.G. (2004). Prednisolone decreases cellular adhesion molecules required for inflammatory cell infiltration in dystrophin-deficient skeletal muscle. Neuromuscul Disord 14, 483–490.

Widrick, J.J., Alexander, M.S., Sanchez, B., Gibbs, D.E., Kawahara, G., Beggs, A.H., and Kunkel, L.M. (2016). Muscle dysfunction in a zebrafish model of Duchenne muscular dystrophy. Physiol Genomics 48, 850–860.

Xi, H., Langerman, J., Sabri, S., Chien, P., Young, C.S., Younesi, S., Hicks, M., Gonzalez, K., Fujiwara, W., Marzi, J., et al. (2020). A Human Skeletal Muscle Atlas Identifies the Trajectories of Stem and Progenitor Cells across Development and from Human Pluripotent Stem Cells. Cell stem cell.

Yang, H.T., Shin, J.H., Hakim, C.H., Pan, X., Terjung, R.L., and Duan, D. (2012). Dystrophin deficiency compromises force production of the extensor carpi ulnaris muscle in the canine model of Duchenne muscular dystrophy. PLoS ONE 7, e44438.

## Supplementary references

Dobin, A., Davis, C.A., Schlesinger, F., Drenkow, J., Zaleski, C., Jha, S., Batut, P., Chaisson, M., and Gingeras, T.R. (2013). STAR: ultrafast universal RNA-seq aligner. Bioinformatics 29, 15–21.

Kuleshov, M.V., Jones, M.R., Rouillard, A.D., Fernandez, N.F., Duan, Q., Wang, Z., Koplev, S., Jenkins, S.L., Jagodnik, K.M., Lachmann, A., et al. (2016). Enrichr: a comprehensive gene set enrichment analysis web server 2016 update. Nucleic Acids Res 44, W90–97.

Liao, Y., Smyth, G.K., and Shi, W. (2014). featureCounts: an efficient general purpose program for assigning sequence reads to genomic features. Bioinformatics 30, 923–930.

Love, M.I., Huber, W., and Anders, S. (2014). Moderated estimation of fold change and dispersion for RNA-seq data with DESeq2. Genome Biol 15, 550.

McCain, M.L., Agarwal, A., Nesmith, H.W., Nesmith, A.P., and Parker, K.K. (2014). Micromolded gelatin hydrogels for extended culture of engineered cardiac tissues. Biomaterials 35, 5462–5471.

Ran, F.A., Hsu, P.D., Wright, J., Agarwala, V., Scott, D.A., and Zhang, F. (2013). Genome engineering using the CRISPR-Cas9 system. Nature protocols 8, 2281–2308.

Schindelin, J., Arganda-Carreras, I., Frise, E., Kaynig, V., Longair, M., Pietzsch, T., Preibisch, S., Rueden, C., Saalfeld, S., Schmid, B., et al. (2012). Fiji: an open-source platform for biological-image analysis. Nature methods 9, 676–682.

Werley, C.A., Chien, M.P., and Cohen, A.E. (2017). Ultrawidefield microscope for high-speed fluorescence imaging and targeted optogenetic stimulation. Biomed Opt Express 8, 5794–5813.

